# Allosteric Modulation of MIF-2 Structure, Catalysis, and Biological Signaling via Cysteine Residues and a Small Molecule, Ebselen

**DOI:** 10.1101/2025.05.15.654344

**Authors:** Vinnie Widjaja, Sirena M. D’Orazio, Pragnya Das, Xander Takada, Divya T. Rajendran, Yuanjun Shi, Iz Varghese, Yannie Lam, Jimin Wang, Victor S. Batista, Vineet Bhandari, George P. Lisi

## Abstract

The macrophage migration inhibitory factor (MIF) family of cytokines comprised of the MIF and D-dopachrome tautomerase (or MIF-2) paralogs share identical tertiary and quaternary structures that contribute to their overlapping enzymatic and signaling activities. Recent investigations of MIF and MIF-2 have shown them to possess N-to-C-terminal allosteric crosstalk, but despite the similarity of this “allosteric pathway,” its regulation of MIF and MIF-2 is not identical. Thus, structure alone does not preserve the precise allosteric mechanism and additional residues that modulate MIF and MIF-2 allosteric function must be characterized. Cysteines have been identified as allosteric switches for the same biochemical functions of MIF and small molecules targeting its N-terminal enzymatic site have affected the structure of three proximal cysteines. Ebselen is a compound that forms covalent selenylsulfide bonds with MIF cysteines and is hypothesized to destabilize and dissociate the MIF trimer into monomers. Ebselen-bound MIF also displays little-to-no catalysis or biological signaling. However, it is unclear whether Ebselen similarly affects the MIF-2 paralog, despite MIF-2 containing two related cysteines (MIF contains three). We used mutagenesis, nuclear magnetic resonance (NMR), molecular dynamics (MD) simulations, *in vitro* and *in vivo* biochemistry to investigate the mechanism of Ebselen as an allosteric modulator of MIF-2 via its cysteines. Our findings suggest that Ebselen partially disrupts the MIF-2 homotrimer, though the overall population of such a structure is <35%, even on the timescale of many hours. Ebselen does attenuate the biological functions of MIF-2 and solution structural biology captures the conformational transitions preceding the destabilized MIF-2 trimer.

**Significance Statement:** The nearly identical MIF and MIF-2 structures have been recently shown to contain an N-to-C-terminal crosstalk between non-overlapping functional sites *in vitro* and *in vivo*. Small molecule inhibitors designed as therapeutics for the MIF superfamily have primarily targeted the N-terminal active site, while very few allosteric molecules have been reported. Ebselen, however, destabilizes MIF by covalently modifying its distal Cys80 residue and disrupting its obligate trimeric assembly. This unique mechanism has never been evaluated in the MIF-2 paralog, and this study reports the structural, dynamic, and functional impact of Ebselen on MIF-2. We define the essential structural hallmarks preceding trimer dissociation and reveal that Ebselen inhibits MIF-2 function before trimer disruption, leading to destabilized monomer-monomer interfaces that slowly degrade the quaternary assembly.

## Introduction

The macrophage migration inhibitory factor (MIF) and D-dopachrome tautomerase (D-DT or MIF-2) cytokines are paralogs that share ~35% sequence identity and a nearly indistinguishable trimeric quaternary assembly (1). This structural similarity is believed to contribute to their overlapping enzymatic activities and interactions with the cluster of differentiation 74 (CD74) receptor. MIF and MIF-2 both play a role in immune cell signaling upon activation of CD74, which include the counter-regulation of glucocorticoids, the migration and recruitment of leukocytes into infectious and inflamed sites, and the sustainment of immune cell survival through inhibition of activation-induced apoptosis (1). Overexpression of both MIF and MIF-2 can lead to chronic inflammatory conditions such as rheumatoid arthritis and asthma. Thus, MIF and MIF-2 appear to toggle between diverse activities, including pro-inflammatory and anti-inflammatory mechanisms.

Previously, the MIF and MIF-2 structures and several conserved functions were found to be regulated by an allosteric coupling of enzymatic activity at the N-terminus to CD74 activation at the C-terminus (2, 3). The allosteric pathway of MIF has been exploited for small molecule inhibition, where ligand binding disrupts the crosstalk between the N- and C-termini (4, 5). Most MIF inhibitors target the N-terminal enzymatic site, though several molecules targeting neither the N-nor C-terminus are also known (4, 6, 7). One such molecule, 2-phenyl-1,2-benzoselenazol-3(2H)-one (Ebselen), was discovered to destabilize MIF through the covalent modification of its Cys80 residue leading to a loss of enzymatic activity and eventual dissociation of the trimer (8, 9). While many examples of small molecule stabilization of nonproductive conformational states have been reported, there has been extremely scant evidence of MIF existing in any form other than its highly stable trimer (10).

Ebselen-induced dissociation has never been explored in the MIF-2 paralog. Despite its similar allosteric pathway and cysteine hotspots, the MIF-2 trimer has a higher thermodynamic stability and slightly different substrate and ligand binding preferences (11). We therefore wonder whether a mechanism of action for Ebselen would translate to MIF-2 (8, 12). Our NMR studies of the MIF-2-Ebselen complex, along with site-directed mutagenesis, confirmed that a single cysteine (Cys23) is modified. This site is distinct from the site of modification within MIF. NMR relaxation experiments suggest Ebselen weakens the monomer-monomer interfaces of MIF-2, despite binding distal to this region. Our biochemical assays reveal that Ebselen attenuates MIF-2 enzymatic activity *in vitro* and CD74 receptor activation *in vivo*, confirming the important regulatory role for cysteine residues in the MIF superfamily. We find the timescale of MIF-2 trimer dissociation to be on the order of days, which is considerably longer than that of MIF. Further, we suggest the mechanism of Ebselen inhibition of MIF-2 to involve allosteric disruption of protein dynamics and multiple biochemical activities as well as dissociation/aggregation of a minor population (<45%) of MIF-2 trimer. Our work provides a further understanding of the allosteric function of MIF-2 and reveals the structural aspects that precede the degradation of its trimeric structure in solution. Additionally, our analysis contributes valuable insights that support ongoing efforts to differentially target the closely related MIF and MIF-2 proteins.

## Results

### The MIF-2 structure surrounding its cysteine residues is impacted by mutations

MIF contains three cysteine residues at positions 56, 59 and 80, only some of which are modified by Ebselen. MIF-2 contains only two cysteines (23 and 56) (8, 12) and to determine the specific sites of Ebselen modification, three MIF-2 variants were designed −C23S, C56S, and C23S/C56S. Prior to studies of these variants with Ebselen, we used ^1^H-^15^N NMR experiments to assess the structural impacts of the mutations themselves. The C23S variant induced a large degree of NMR chemical shift perturbations (CSPs), while those of C56S were comparatively muted (**Fig. S1**). These data suggest that Cys23 is the more critical allosteric structural handle in MIF-2. The strongest CSPs (largest green spheres in **Fig. S1**) surrounded the site of mutation and Cys56, the other Cys residue. This observation suggests a manner of crosstalk between the Cys sites, despite their spatial separation. Line broadening of NMR resonances (**Fig. S1**, blue spheres) occurs at sites primarily localized to the C-terminus. Overall, Cys23 appears more critical to MIF-2 structure than Cys56, and a C23S/C56S double mutant displays a mostly additive effect on CSPs (**Fig. S1**). Despite these local atomistic effects, the overall fold of MIF-2 remains the same, as does its secondary structure (**Fig. S2**). Thermal stabilities of the C56S and C23S/C56S variants are diminished, while WT and C23S have similar thermal stabilities, which is surprising given the apparent structural importance of Cys23 to MIF-2.

### Ebselen selectively modifies Cys23 of MIF-2

In previous studies of MIF, it was found that Cys80 was the major site of modification by Ebselen (8, 12). Like Cys80 of MIF, Cys23 of MIF-2 is located on a solvent-exposed helix, leading us to hypothesize that Cys23 would be modified by Ebselen after incubation with WT MIF-2 and MIF-2 variants for 1 hour prior to NMR data collection. To assist with solubility of Ebselen and minimize NMR spectral artifacts of Ebselen precipitation during titration with MIF-2, a background of 15% (v/v) DMSO-D_6_ was used without significantly altering MIF-2 structure (13). ^1^H-^15^N transverse relaxation optimized spectroscopy-heteronuclear single quantum coherence (TROSY-HSQC) spectral overlays of apo MIF-2 (WT and variants) and MIF-2 bound to Ebselen revealed that Ebselen induces NMR CSPs in WT MIF-2 and C56S (**Fig. 1A** and **Fig S3**), suggesting that Cys23 is the site of modification. Ebselen does not seem to interact with C23S and (as expected) C23S/C56S, evidenced by a lack of CSPs. The Ebselen-induced CSP profiles of WT MIF-2 and C23S are very similar, localizing around Cys23 with many residues sharing the same shift trajectories, suggesting similar ligand bound conformations (**Fig. 1B, C**). Regions adjacent to the α-helix 2 that houses Cys23 have also broadened, implying a change in its dynamics.

**Figure 1.**
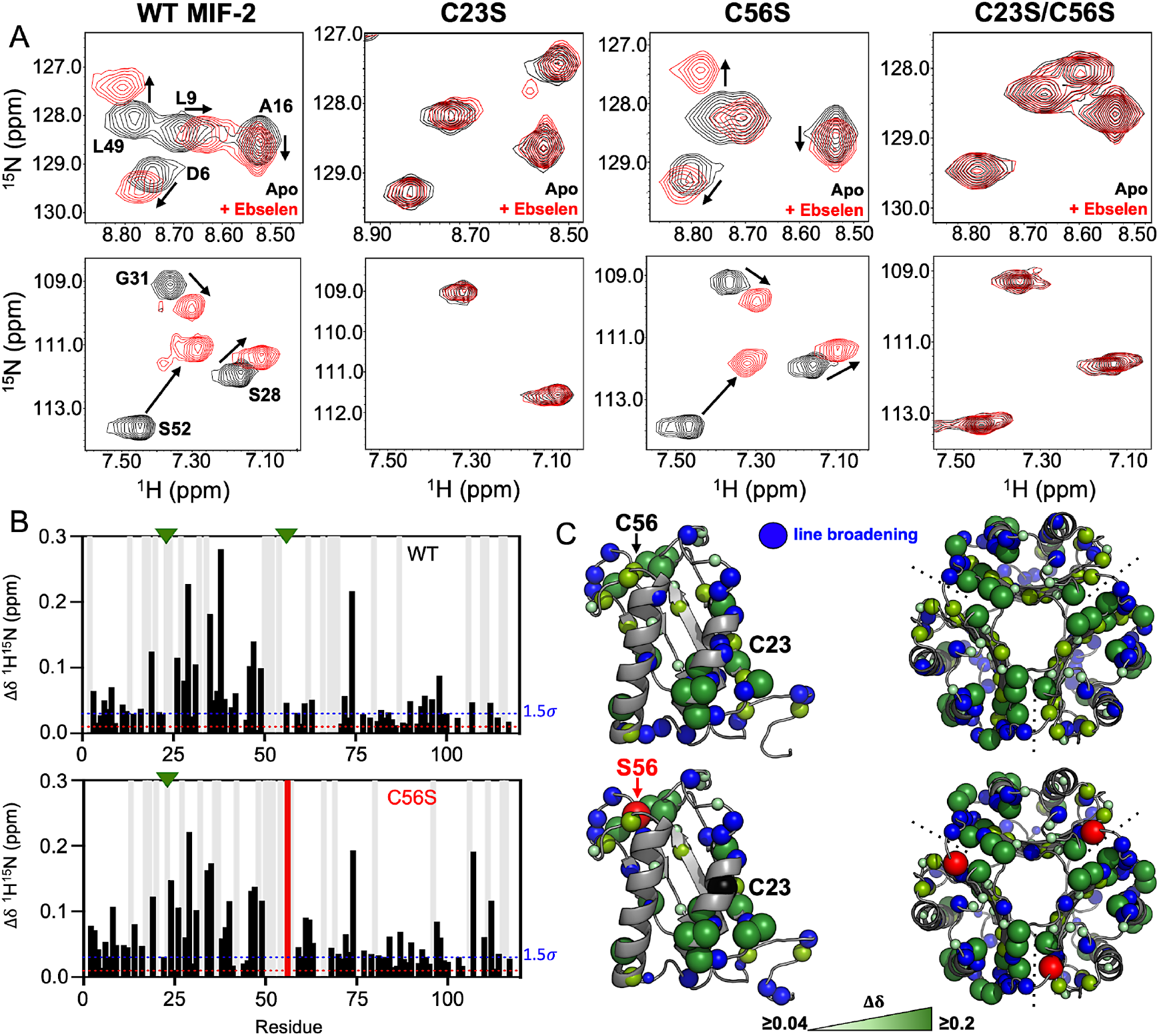
Structural effect of Ebselen on WT MIF-2 and MIF-2 variants. **(A)** Representative snapshots of ^1^H-^15^N HSQC spectral overlays of apo (black) and Ebselen-bound (red) MIF-2. NMR CSPs suggest Ebselen selectively modifies Cys23 of WT MIF-2 and the C56S variant but does not interact with C23S (via Cys56) or C23S/C56S MIF-2. **(B)** Per-residue NMR CSPs (black bars) caused by Ebselen modification of WT and C56S MIF-2. Light gray bars denote sites of NMR line broadening. Red bars denote a Cys-to-Ser mutation, and green triangles denote a native Cys residue. **(C)** CSPs >1.5σ of the 10% trimmed mean of all shifts are mapped on the MIF-2 monomer (green spheres). Sphere size correlates with the intensity of the CSP. Blue spheres represent sites of NMR line broadening, red spheres denote a Cys-to-Ser mutation, and black spheres denote a native Cys residue.

Previously, the reduction, alkylation, or blocking of MIF cysteines by chemical cross-linking was shown to prevent Ebselen-induced MIF aggregation (8, 13). To test a similar effect in MIF-2, we reduced its cysteines with the addition of 5 mM DTT to the NMR tube. We observed no Ebselen-induced change in the WT MIF-2 NMR spectrum or precipitation (**Fig. S4**). Thus, Ebselen modification of MIF and MIF-2 appears to rely on redox-neutral or oxidizing conditions to modify the protein.

### MD simulations highlight the specific contacts within MIF-2 that stabilize Ebselen

After localizing the specific interaction point of Ebselen with MIF-2, we next investigated the molecular contacts facilitating the complex. MD simulations generated an equilibrated structure of the MIF-2-Ebselen complex, revealing a network of hydrophobic contacts surrounding the Cys23 site (**Fig. 2A, B**), facilitated by the aromatic rings of the Ebselen molecule. Aliphatic residues Ala24, Ala27, Ala34, Ala48, and Gly51 (backbone) comprise the major contacts, while van der Waals interactions are observed with the polar Lys20 and Thr53 residues. A comparison of unliganded MIF-2 and MIF-2-Ebselen structures reveals that Ebselen binding alters the architecture of the Cys23 pocket, especially the Ala35 and Gly51 backbone regions and the Lys20 side chain (**Fig. 2C**). Importantly, NMR reveals a chemical shift perturbation or line broadening event at each of the residues shown to make contacts with Ebselen *in silico*, supporting the binding pose (**Fig. 2D**). Analysis of the root-mean-square fluctuations (RMSF) surrounding the Ebselen binding site in each MIF-2 monomer reveals subtle changes to the protein dynamics (**Fig. S5**, ΔRMSF ≤ 0.5 Å). Overall, the individual monomers of MIF-2 behave very similarly in response to Ebselen. An Ebselen-induced enhancement of MIF-2 motions is visible at the Cys23 binding site and the surrounding region, consistent with NMR line broadening (**Fig. 2D**). A similarly small, but noticeable, enhancement of MIF-2 RMSF occurs near Gly51, which is also line broadened in NMR experiments.

**Figure 2.**
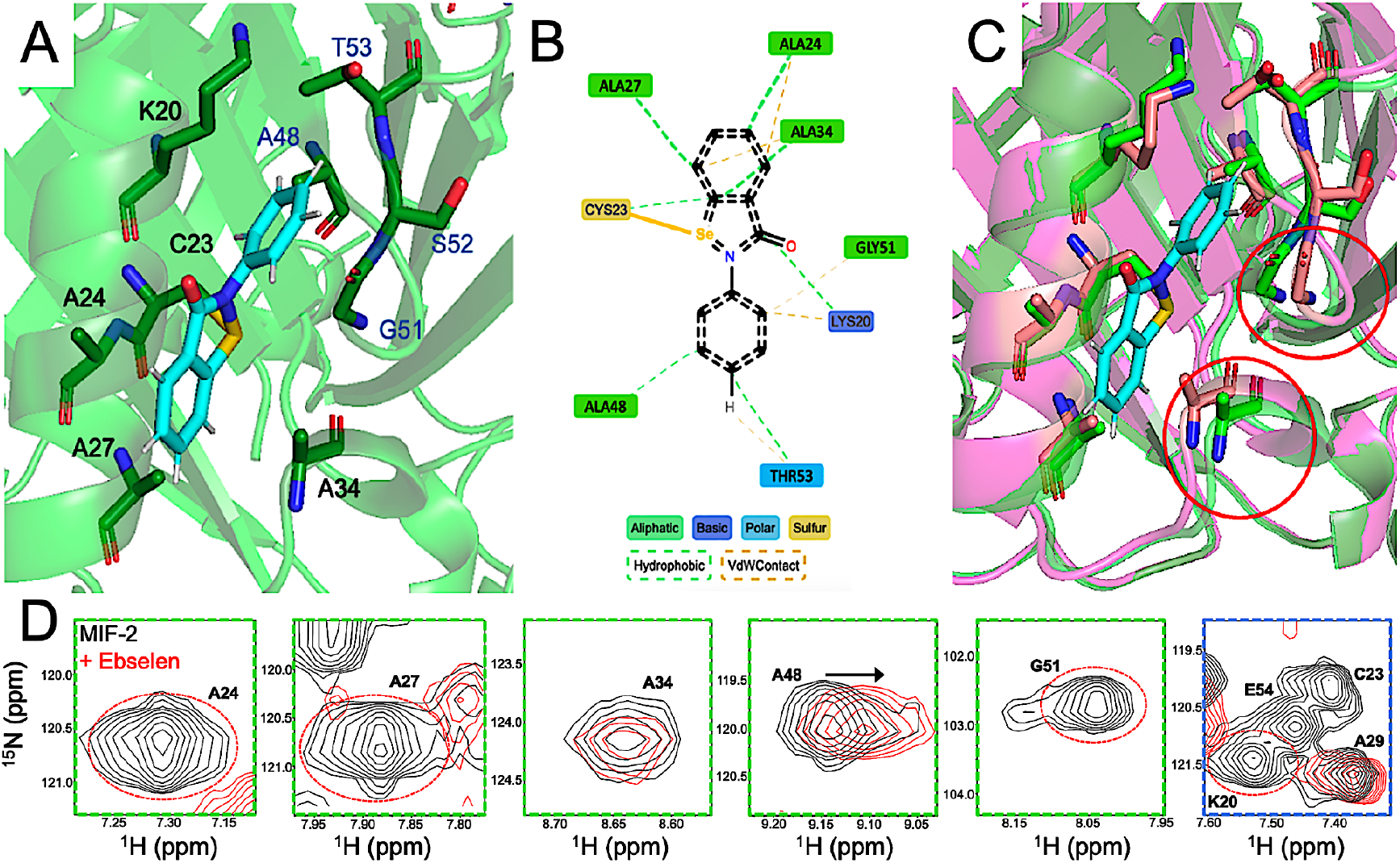
Molecular interactions between MIF-2 and Ebselen. **(A)** Equilibrated structure of the MIF-2-Ebselen complex from MD simulations. Residues interacting with Ebselen are shown in sticks. **(B)** Cartoon view of the MIF-2 interactions with Ebselen. **(C)** Alignment of the Ebselen-bound structure (green) with the structure of apo MIF-2 (magenta). Major structural differences are highlighted with red circles. **(D)** NMR suggests an interaction between MIF-2 and Ebselen at each of the residues highlighted in **(B)**, using the same color-coded legend. A48 shows a clear CSP, while other resonances are broadened (most beyond detection, red dashed circles) in the Ebselen-bound state.

### Ebselen perturbs local dynamics of MIF-2, but does not disrupt the trimer on the NMR timescale

NMR spin relaxation experiments are highly sensitive to multi-timescale conformational dynamics that influence protein function and ligand binding (14). These include pico-nanosecond bond vector fluctuations that organize binding sites and report on global tumbling (15). We previously showed the degree of motion on this timescale to be predictive of MIF and MIF-2 catalytic function and reflect the stability of the monomer-monomer interfaces (2, 3). To determine how Ebselen modification of MIF-2 affects its intrinsic dynamics and global tumbling, we used NMR spin relaxation. *T*_1_ and *T*_2_ values measured for WT MIF-2 were 1154 ± 51 ms and 51 ± 5 ms, respectively, consistent with prior reports (16). The C56S variant showed similar structural compactness, with *T*_1_ and *T*_2_ values of 1020 ± 42 ms and 56 ± 3 ms, respectively (**Table 1**). When bound to Ebselen, *T*_1_ and *T*_2_ values for WT MIF-2 and C56S were 1142 ± 59 ms and 55 ± 4 ms and 1094 ± 55 ms and 53 ± 4 ms, respectively (**Table 1**). These data indicate that Ebselen modification did not significantly alter the global tumbling of MIF-2. We estimated the rotational correlation times (τc) from these data (17), confirming that τc for Ebselen bound forms of MIF-2 are very similar to τc of the apo proteins (**Table 1**). The calculated τc values are also in line with a compact protein of ~35 kDa (16) and importantly, suggest that Ebselen does not dissociate the MIF-2 trimer on the timescale of the NMR experiment.

**Table 1.** Effect of Ebselen on the MIF-2 molecular weight and oligomeric state. Residue-averaged *T*_1_ and *T*_2_ values for WT MIF-2 and C56S MIF-2 in the absence and presence of Ebselen are nearly identical. Estimated rotational correlation times (τ_c_) for each state are similar for each state and consistent with a ~35 kDa protein.

| WT MIF-2 |  | C56S MIF-2 |  |
| --- | --- | --- | --- |
| $\langle T_1 \rangle$ (ms) | $1154 \pm 51$ | $\langle T_1 \rangle$ (ms) | $1020 \pm 42$ |
| $\langle T_2 \rangle$ (ms) | $51 \pm 5$ | $\langle T_2 \rangle$ (ms) | $56 \pm 3$ |
| $\tau_c$ (ns) | $15 \pm 0.4$ | $\tau_c$ (ns) | $13 \pm 0.4$ |
| WT MIF-2 + Ebselen |  | C56S MIF-2 + Ebselen |  |
| $\langle T_1 \rangle$ (ms) | $1142 \pm 59$ | $\langle T_1 \rangle$ (ms) | $1094 \pm 55$ |
| $\langle T_2 \rangle$ (ms) | $55 \pm 4$ | $\langle T_2 \rangle$ (ms) | $53 \pm 4$ |
| $\tau_c$ (ns) | $15 \pm 0.7$ | $\tau_c$ (ns) | $14 \pm 0.5$ |

Though average relaxation parameters of MIF-2 appear unaffected by mutation or by Ebselen, per-residue analysis highlights differences in flexibility of specific regions of the protein as a result of Ebselen modification. We hypothesized that the greatest perturbations to protein dynamics would occur near the monomer-monomer interface, where Ebselen has been reported to destabilize MIF (8, 12). Based on rotational correlation times (suggesting an intact trimer), dissection of residue-specific MIF-2 flexibility will reveal the changes that precede trimer dissociation, providing a window into the conformational reshuffling on the fast timescale. Correlation plots comparing *R*_2_/*R*_1_ relaxation rates for apo proteins (*i*.*e*. WT MIF-2 or C56S) and the respective Ebselen bound states show many residues to be affected by Ebselen (**Fig. 3A, C**). Residues with *R*_2_/*R*_1_ parameters outside of the red dashed boundaries (+1.5σ of the 10% trimmed mean) of each plot denote significant Ebselen-induced change in fast timescale flexibility. When mapped onto the MIF-2 structure (**Fig. 3B, D**), these effects surround the Cys residues, particularly Cys23 on α-helix 2 that is adjacent to the monomer-monomer interface. This location is consistent with NMR CSPs and line broadening that qualitatively signifies heightened dynamics (**Figs. S1** and **Fig. 1**). The observed dynamic effects are, in this case, most strongly influenced by *R*_2_. Thus, regions of orange highlighted on the MIF-2 monomer in **Fig. 3** indicate elevated *R*_2_ values of the Ebselen bound state (*i*.*e*. enhanced flexibility), while blue regions were sites where apo protein dynamics were suppressed by Ebselen. A further level of detail was obtained by examining the per-residue *R*_1_ and *R*_2_ values (**Fig. S6**). Here, the effect of Ebselen was subtle, except for a notable change in *R*_2_ occurring adjacent to the Cys23 binding site. Comparison of the WT and C56S proteins (**Fig. S6**) highlights a distinct effect of Ebselen on the C56S C-terminus, spanning residues I107 – T115, which also have elevated *R*_2_ values.

**Figure 3.**
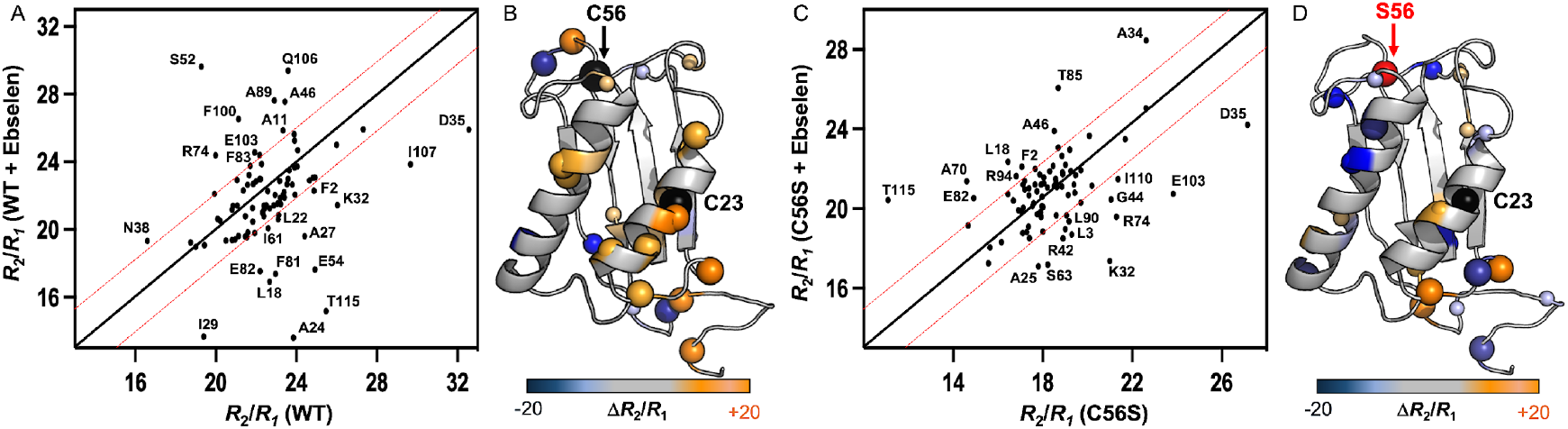
Effect of Ebselen on MIF-2 dynamics. **(A)** Correlation plot of *R*_2_/*R*_1_ relaxation rates for WT MIF-2 and WT MIF-2 + Ebselen. Points falling outside the red dashed boundaries (+1.5σ of the 10% trimmed mean) are sites of significant dynamic perturbation in the presence of Ebselen, which are mapped onto MIF-2 in **(B)** as the magnitude of Δ*R*_2_/*R*_1_ (Ebselen – Apo). **(C)** Correlation plot of *R*_2_/*R*_1_ relaxation rates for C56S MIF-2 and C56S MIF-2 + Ebselen. As in **(A)**, points outside the red dashed boundaries are sites of significant dynamic perturbation in the presence of Ebselen, which are mapped onto MIF-2 in **(D)** as the magnitude of Δ*R*_2_/*R*_1_ (Ebselen – Apo).

### Solvent PRE confirms an effect of Ebselen at the MIF-2 monomer-monomer interface

To investigate whether the dynamics of trimer dissociation can be captured by other metrics, we used solvent paramagnetic relaxation enhancement (sPRE). The sPRE effect across the protein sequence reports on areas that are solvent accessible due to line broadening caused by the TEMPOL nitroxide radical (18). When examining MIF-2 alone, a sPRE effect is only observed in regions that are solvent exposed in crystal structures (**Fig. 4A, B**). When MIF-2 is bound to Ebselen, the sPRE effect is dramatically enhanced, with regions of solvent exposure again surrounding Cys23, with the monomer-monomer interface showing the largest decreases in resonance intensity (**Fig. 4B-D**). We note, however, that residual peak intensities (*I*_*WT*_ – *I*_*Ebselen*_) are strongly positive across the entire MIF-2 sequence (**Fig. 4C**), suggesting a widespread deprotection of MIF-2 from solvent. Collectively, solution NMR indicates that MIF-2 remains a trimer in the NMR tube, but that its monomer-monomer interfaces are destabilized, allowing for increased solvent exposure of the MIF-2 core.

**Figure 4.**
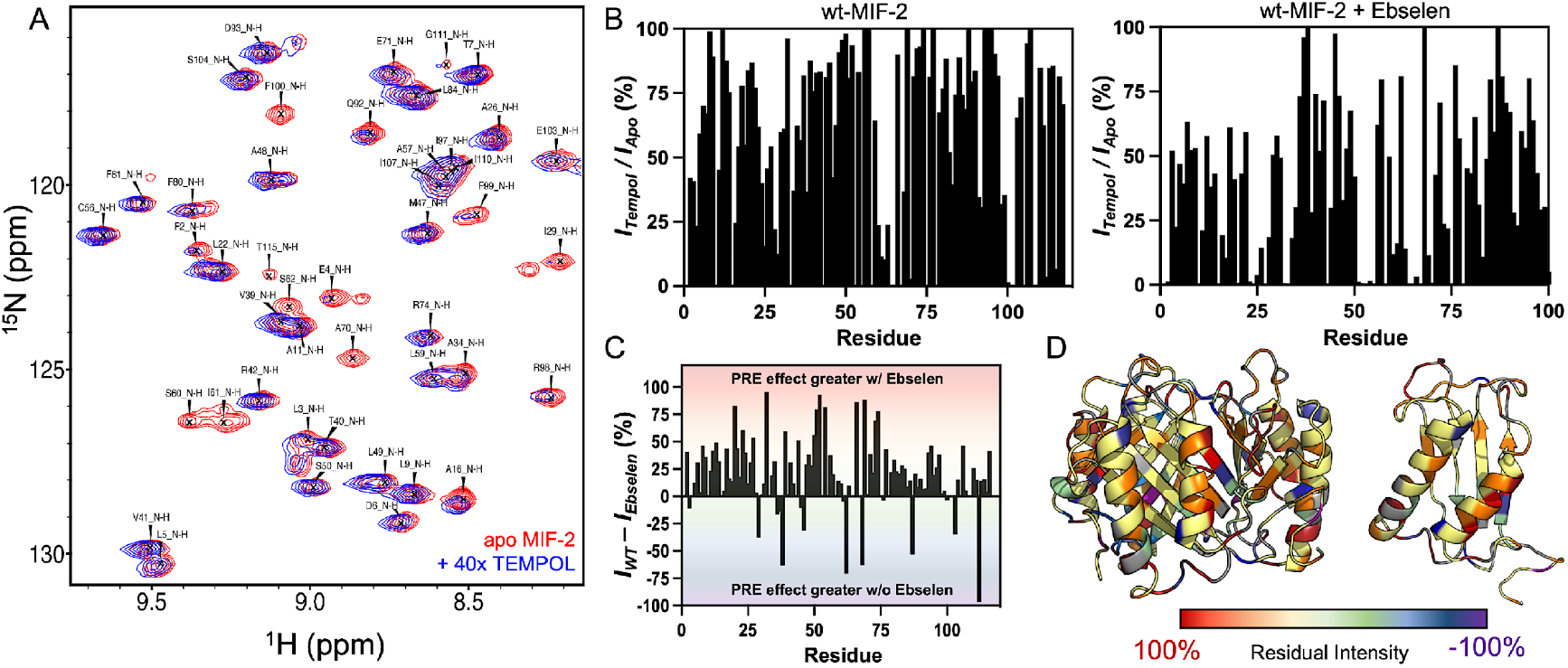
Effect of Ebselen on solvent accessibility at the MIF-2 monomer interface. **(A)** Snapshot of a solvent PRE experiment showing the specific disappearance of resonances from the WT MIF-2 NMR spectrum when TEMPOL is present in the buffer (blue, while the WT MIF-2 reference spectrum is in red). **(B)** Intensity ratios (*I*_*tempol*_/*I*_*apo*_) describing the per-residue effect of TEMPOL on apo WT MIF-2 (left) and WT MIF-2 bound to Ebselen (right). **(C)** Difference plot (*I*_*WT*_ - *I*_*Ebselen*_) highlighting the substantially larger PRE effect on Ebselen-bound MIF-2. **(D)** Residual intensity calculated in **(C)** is mapped onto the MIF-2 trimer and monomer. The strongest PRE effect is shown for Ebselen-bound MIF-2 at the monomer interfaces (warm colors).

### Ebselen allosterically impacts MIF-2 biochemistry and slowly degrades the trimer

Concentrations of Ebselen (15 uM) were previously shown to inhibit MIF tautomerase activity (8). MIF-2 contains the same tautomerase active site with a conserved catalytic Pro1 at its N-terminus (19). This site has been exploited as a drug discovery target, as enzymatic inhibition was shown to affect MIF and MIF-2 signaling via CD74 (19, 20). We measured the enzymatic activity of MIF-2 and found that Ebselen diminished this function by ~45% (**Fig. 5A**). We attribute the decrease in tautomerase activity to the structural changes surrounding Cys23 and the monomer interfaces that comprise the active site. Ebselen attenuated the tautomerase activity of C56S MIF-2 to nearly the same degree, consistent with Cys23 being the site of Ebselen modification. Thus, Cys23 is a novel allosteric node for MIF-2 catalytic function that was uncharacterized in prior work.

**Figure 5:**
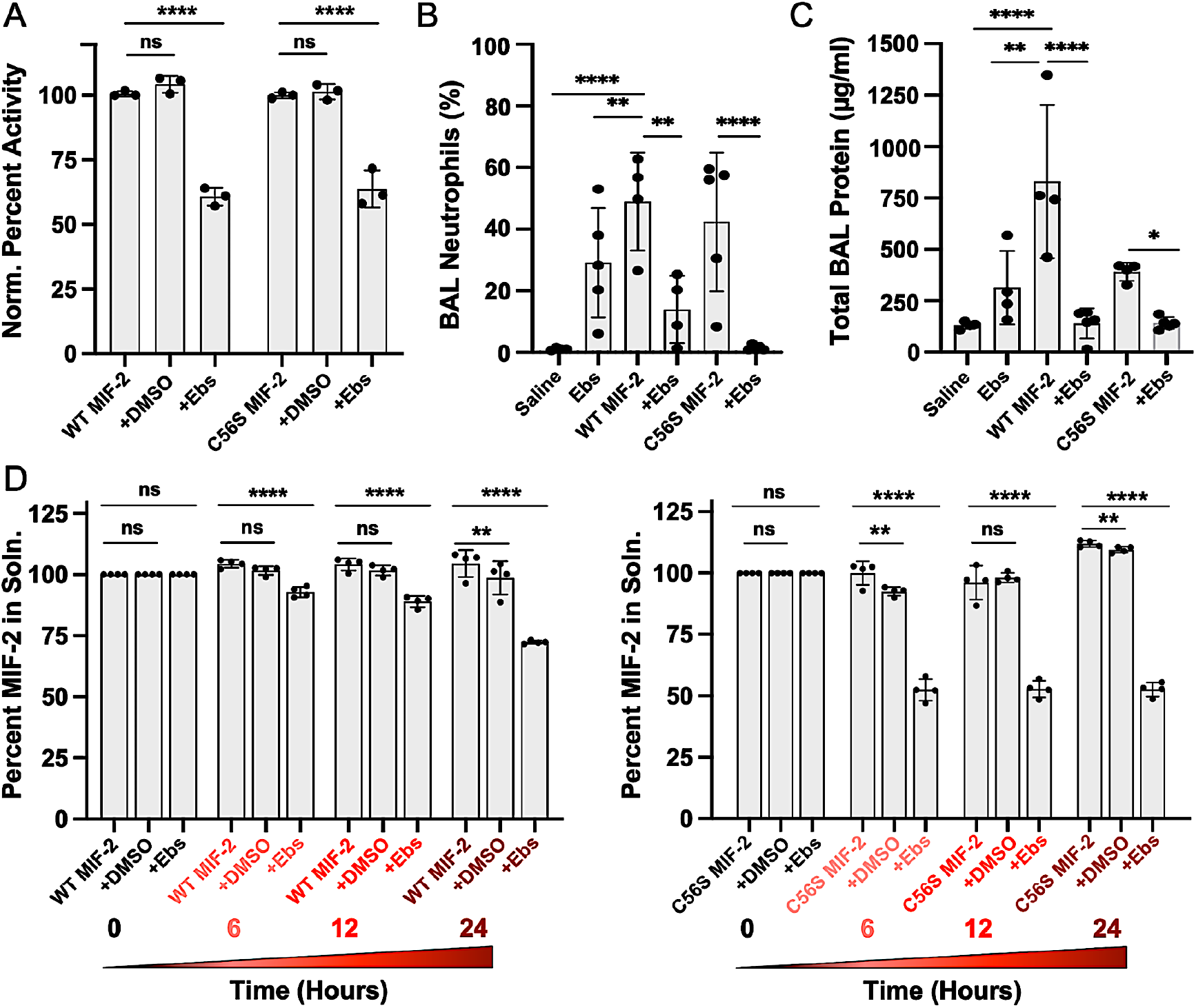
Effect of Ebselen on the function of MIF-2. **(A)** Enzymatic tautomerase assays with a 4-HPP substrate show full activity for WT MIF-2 and WT MIF-2 in the presence of the equivalent volume of DMSO used to solubilize Ebselen (control). Incubation of MIF-2 with Ebselen attenuates enzymatic activity. A similar trend is observed for C56S MIF-2. Data are reported as mean ± SEM (n = 3 for each group). ****p<0.0001 **(B)** Ebselen-bound MIF-2 attenuates neutrophil recruitment in murine BAL fluid *in vivo* compared to MIF-2 alone, Ebselen alone, or saline control. A similar trend is observed for C56S MIF-2. Data are expressed as mean ± SEM (n = 4-5 for each group). **p<0.005; **\*\*\*\***p<0.0001 **(C)** Ebselen-bound MIF-2 attenuates BAL fluid protein levels *in vivo* (a surrogate marker for alveolar-capillary leak and pulmonary edema) compared to MIF-2 alone, Ebselen alone, or saline control. A similar trend is observed for C56S MIF-2. Data are expressed as mean ± SEM (n = 4 for each group). *p<0.05; **p<0.005; **\*\*\*\***p<0.0001 **(D)** Spin-down assay quantifying the percent of soluble WT or C56S MIF-2 in solution over a 24-hour period (light-to-dark red colors) using MIF-2 alone, MIF-2 with the equivalent volume of DMSO used to solubilize Ebselen (control), and MIF-2 with Ebselen. Data are reported as mean ± SEM (n = 4 for each group). **p<0.005; **\*\*\*\***p<0.0001

We next assessed the effect of Ebselen on the ability of MIF-2 to activate CD74 *in vivo*, via neutrophil recruitment to murine lungs (21–23). Following the intratracheal delivery of saline control, Ebselen control, WT MIF-2 (± Ebselen), or the C56S variant (± Ebselen), the percentage of neutrophils and total protein (a marker for pulmonary edema) were quantified from the bronchoalveolar lavage (BAL) fluid (**Fig. 5B, C** and **Fig. S7**). As expected, WT MIF-2 stimulated CD74-dependent neutrophil recruitment and increased the total BAL protein levels, relative to control cases. Ebselen-bound MIF-2 showed significantly attenuated neutrophil recruitment and BAL protein, consistent with an inhibitory effect. The C56S variant displayed comparable neutrophil recruitment activity to that of WT MIF-2, reflecting the modest structural impact of this mutation on MIF-2. A complex with Ebselen again suppresses neutrophil recruitment and BAL protein levels. Collectively, these data suggest that the structural and dynamic impacts of Ebselen binding propagate from the distal Cys23 to the MIF-2 C-terminus. Allosteric control of CD74 binding now appears to involve C-terminal residues beyond those studied here, since only one mutation, T112A, suppresses CD74 activation (3).

Though NMR suggested the soluble MIF-2-Ebselen complex to remain trimeric, we attempted to quantify any population of MIF-2 found in a precipitate. We used a time-dependent spin down assay to separate any visible precipitate, which has been shown to contain unstable monomers (8, 12), from the MIF-2-Ebselen solution (**Fig. 5D**). Over a period of 24 hours, nearly 70% of MIF-2 remains in solution, while ~30% is spun out as precipitate. The dissociation of MIF-2 therefore happens quite slowly, with more than 12 hours required to reduce the soluble fraction of MIF-2 by even 20%. Importantly, our assays point specifically to Ebselen as a driver of solubility of MIF-2, as a control time course with the equivalent volume of DMSO required to solubilize Ebselen has little-to-no effect of MIF-2. The same spin down assay conducted on C56S MIF-2 revealed an identical trend, albeit with a greater population of C56S (~50%) found in the precipitate (**Fig. 5D**). Here, Ebselen-induced destabilization of the trimer occurs after 6 hours and remains stable beyond that point. We attribute the apparent enhancement of the Ebselen effect to the notable loss of thermostability of the C56S MIF-2 trimer, compared to WT MIF-2 (**Fig. S1**).

## Discussion

Molecular crosstalk between the N-terminal enzymatic and C-terminal CD74 activation sites of MIF and MIF-2 are conserved features of the trimeric assemblies, despite some subtle variations in amino acid composition (2, 3). Recent work has expanded the allosteric network of MIF to include distal cysteine residues that toggle the MIF structure and fine tune functional activity (24–26). The structural similarity between MIF and MIF-2 has led to hypotheses of synergistic or compensatory functions (27), thus it is important to understand the molecular details of the MIF-2 paralog. Explorations of anti-inflammatory or anti-tumor therapies (28, 29) targeting MIF identified Ebselen, a multifunctional small molecule that downregulates inflammation and protects cells from oxidative damage (29, 30). Ebselen has also been used as a cysteine-modifying drug in studies of redox mechanisms and protein aggregation in amyotrophic lateral sclerosis (ALS), Parkinson’s disease, and cerebral ischemia (29, 31, 32).

We used Ebselen as a probe to evaluate the allosteric impact of MIF-2 cysteine residues and found that Ebselen covalently modifies Cys23 of MIF-2, located on a solvent exposed α-helix opposite the site of modification (Cys80) in MIF (8, 12). Site-directed mutagenesis confirms the point of modification, but also independently revealed the MIF-2 cysteines as allosteric nodes that altered its structure and function. NMR revealed Ebselen-bound MIF-2 remains trimeric in solution, albeit with a distorted monomer-monomer interface, and biochemistry provided evidence that trimer destabilization and MIF-2 aggregation occurs on a timescale of several hours-to-days (**Fig. 6**). However, the portion of MIF-2 rendered as an unstable precipitate (and presumably monomeric) by Ebselen represents a minor fraction (<50%), while the majority remains in solution with diminished biochemical and biological activities.

**Figure 6.**
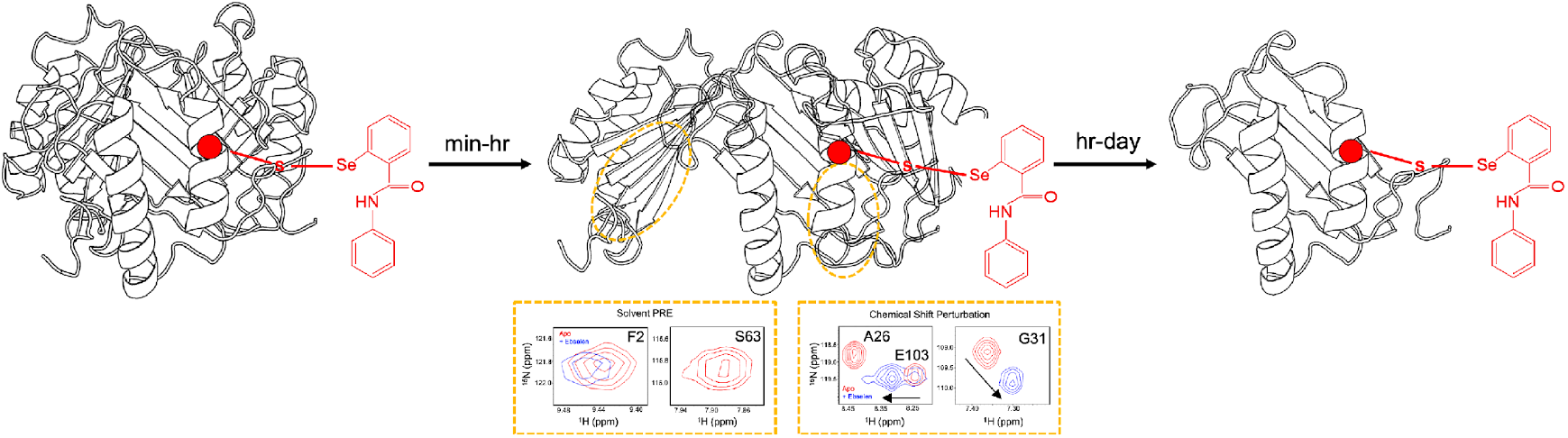
Cartoon scheme summarizing the effect of Ebselen on MIF-2. Modification of accessible Cys23 by Ebselen (red molecule) induces structural and dynamic changes at the monomer-monomer interfaces, observed by NMR CSPs and solvent PRE (yellow dashed circles). At longer time points, visible protein precipitation is apparent, consistent with a disruption of the trimer structure and formation of disordered and aggregation-prone monomers.

Ebselen has furthered an appreciation for the critical influence of cysteine residues on the allosterically coupled N-terminal catalytic tautomerase activity and C-terminal CD74 activation of the MIF superfamily (2, 3, 33, 34). Prior work reported Cys56, Cys59, and Cys80 of MIF to each play distinct roles in its redox chemistry and allosteric control. Conformational dynamics surrounding the C_56_-A-L-C_59_ thioredoxin-like motif were shown to be critical for therapeutic antibody targeting of MIF (35, 36), as were redox-dependent modifications of each site (37). Mutation or modification of Cys80 was also shown to shift the MIF structural equilibrium toward a latent conformer that, when disrupted, abolished CD74 activation (25). The cysteine residues of MIF-2 have never been explored as allosteric modulators of function, partially because only Cys56 of MIF-2 is conserved between the proteins. Cys23, like Cys80 of MIF, is located on a solvent exposed α-helix, though on the opposite side of the monomer. However, like Cys80, our biochemical studies show that mutation of Cys23 diminishes catalytic function, CD74 activation and inflammation in murine lungs. When modified with Ebselen, similar effects are observed, highlighting its contribution to the allosteric network of MIF-2.

It is important to note that subtle differences between the MIF and MIF-2 trimers at the biophysical level may differentiate their biological roles. It was recently suggested that CD74 binding and activation may be dictated by a small stretch of C-terminal residues, and many chemically similar small molecules show a preference for one paralog over another, despite their identical tertiary structures (38–40). Prior studies of the MIF-Ebselen complex showed an almost total dissociation of the MIF trimer over the course of one hour (8, 12) while our work with MIF-2 suggests ~35% of the total Ebselen-bound protein aggregates and precipitates after 24 hours. Nonetheless, Ebselen remains the only known compound capable of disrupting this stable (~micromolar affinity) (8, 12) and compact trimeric structure, which is widely considered to be the biologically active form.

## Conclusion

We report the cysteine residues of MIF-2 as previously unrecognized allosteric handles that, when modified by the small molecule Ebselen, undergo local structural changes that propagate to the interfaces of its trimer structure to disrupt its enzymatic and cellular signaling functions. Cysteine-mediated allostery within the well-studied MIF-2 paralog, MIF, was confirmed only in the past few years. Despite Ebselen-induced destabilization, a substantial portion of the MIF-2 trimer remains intact in solution, suggesting greater structural resilience compared to MIF. NMR highlights the atomic level structural changes that prime the MIF-2 trimer for eventual dissociation which given the widely hypothesized pathophysiological roles of the MIF superfamily, opens new avenues for therapeutic intervention.

## Materials and Methods

### Protein Expression and Purification

Wild-type and mutant MIF-2 was expressed and purified as previously described (41). To produce unlabeled protein for biochemical assays, a pET-22b plasmid encoding MIF-2 was transformed into BL21-Gold (DE3) *E. coli* cells and grown in Luria-Bertani (LB) medium at 37°C to an OD_600_ 0.6-0.8. Protein expression was induced with 1 mM IPTG followed by shaking at 20°C for 16 hrs. Cells were harvested by centrifugation and resuspended in a buffer of 20 mM Tris and 20 mM NaCl at pH 8.5, supplemented with 1 mM phenylmethylsulfonyl fluoride. Cells were lysed by sonication on ice and then centrifuged to remove cell debris. The supernatant was filtered through a 0.22 *μ*m filter, loaded onto a Q-Sepharose (anion-exchange) column, washed with a buffer of 20 mM Tris and 20 mM NaCl at pH 8.5, and eluted with 5% of a buffer of 20 mM Tris, and 1 M NaCl at pH 8.5. Additional contaminants were removed by size-exclusion chromatography with a HiLoad 16/600 Superdex 75 column. Protein purity was assessed by sodium dodecyl-sulfate polyacrylamide gel electrophoresis (SDS-PAGE) and the concentration of MIF-2 (expressed per monomer) was determined using ε_280_ = 5500 M^−1^cm^−1^ and a Pierce Bicinchoninic Acid Protein Assay Kit (Thermo Fisher Scientific).

### NMR Spectroscopy

Isotopically labeled NMR samples of MIF-2 were prepared as described earlier, but rather than LB medium, MIF-2 expression was carried out in M9 minimal media containing MEM vitamins (Sigma-Aldrich) and ^15^NH_4_Cl (Cambridge Isotope Labs) as the sole nitrogen source. Purified MIF-2 was dialyzed into a final NMR buffer of 20 mM Tris, 20 mM NaCl at pH 7.4 with 10% D_2_O and then concentrated to 0.5-1.0 mM. NMR experiments were performed on Bruker Avance NEO 600 MHz spectrometer at 30 °C. NMR data were processed using NMRPipe and analyzed in Sparky along with in-house scripts (42, 43).

### Ebselen titrations of ^15^N-MIF-2

^1^H-^15^N TROSY HSQC fingerprint spectra of 0.5 mM MIF-2 were collected at 30 °C in 20 mM Tris and 20 mM NaCl at pH 7.4 with 10% D_2_O. Ebselen (Focus Biomolecules) was dissolved in 100% DMSO-d_6_ at a stock concentration of 200 mM. Titrations were performed by adding small aliquots of Ebselen to the MIF-2 sample with gentle mixing by pipette and a 30-minute incubation time before the acquisition of spectra. Saturation of MIF-2 was followed until NMR chemical shift perturbations were no longer visible with subsequent additions of Ebselen. NMR chemical shift perturbations were quantified using the method of Bax and coworkers (44). *NMR*

### Spin Relaxation

TROSY-based spin relaxation experiments were performed with the ^1^H and ^15^N carriers set to the water resonance and 120 ppm, respectively. Longitudinal relaxation rates (*R*_*1*_) were measured with randomized *T*_1_ delays of 0, 20, 60, 100, 200, 600, 800, 1,200, 1,500, 2,000, and 2500 ms. Transverse relaxation rates (*R*_*2*_) were measured with randomized *T*_*2*_ delays of 0, 16.9, 33.9, 67.8, 136, 169, and 203 ms. Relaxation data were collected in a temperature-compensated manner with eight scans of 1024 and 256 points in the direct and indirect dimensions, respectively, over a 14 ppm ^1^H and 35 ppm ^15^N spectral width. The recycle delay in these experiments was 2.5 s (45). Longitudinal and transverse relaxation rates were extracted by nonlinear least squares fitting of the peak heights to a single exponential decay using in-house software. Uncertainties in these rates were determined from replicate spectra with duplicate relaxation delays of 20 (×2), 60, 200, 600 (×2), 800, and 1200 ms for *T*_1_ and 16.9, 33.9 (×2), 67.8, 136 (×2) and 203 ms for *T*_2_. The values of rotational correlation times (τ_*c*_) were estimated using the following equation, where *v*_*N*_ is the ^15^N frequency in Hertz (46):

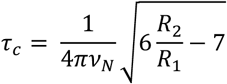

### Solvent PRE

^1^H-^15^N TROSY HSQC spectra of 0.5 mM MIF-2 were collected at 30 °C in a buffer of 20 mM Tris and 20 mM NaCl at pH 7.4 with 10% D_2_O. 4-Hydroxy-2,2,6,6-tetramethylpiperidine-1-oxyl (TEMPOL, Fisher Scientific) was dissolved in the same buffer at a stock concentration of 5 M. PRE titrations were carried out by adding increasing amounts of TEMPOL to the MIF-2 NMR sample to a final ratio of 1:40 (MIF-2:TEMPOL), reflecting the maximum decrease in NMR peak intensity. Ebselen was added to an identical NMR sample of MIF-2, incubated at room temperature for 1 hour, and TEMPOL was again titrated to a ratio of 1:40 (MIF-2: TEMPOL). Data were analyzed based on the relationship of TMEPOL-saturated MIF-2 versus MIF-2 alone (MIF-2_tempol_/ MIF-2_apo_ or MIF-2 + Ebselen_tempol_/MIF-2 + Ebselen_apo_).

### Circular Dichroism Spectroscopy

Circular dichroism data were collected on a Jasco J-815 spectropolarimeter using a 0.2-cm quartz cuvette with 10 μm MIF-2 in a buffer of 20 mM sodium phosphate at pH 7.4. Thermal denaturation experiments were collected at 218 nm over a temperature range of 25 to 90 °C, sampling 1.5 °C at a rate of 1.5 °C/min. Thermal unfolding profiles were fit to the following equation in GraphPad Prism:

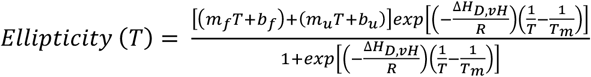

### Molecular Dynamics Simulations

The 1.27 Å resolution crystal structure of human MIF-2 (PDB ID: 7MSE) was aligned to the 2.60 Å MIF structure (PDB ID: 1MIF) and used as the receptor for docking. Ebselen was docked to MIF-2 using AutoDock Vina (47) via UCSF Chimera (48). Among the top 10 predicted binding poses, the one closest to Cys23 - the proposed binding site - was selected as the representative binding pose for subsequent molecular dynamics (MD) simulations. Ebselen parameters and topology were obtained from the Automated Topology Builder (ATB) repository (49) and converted to AMBER format. MIF-2 parameters were generated using the tLEaP program from the AmberTools suite (50, 51). Initial preparation of the MIF-2 structure for MD simulations was performed using Schrödinger Maestro (52) - the protein termini were capped, protonation states were predicted with PropKa (53), and the structure was minimized using the OPLS force field (54) with a 0.3 Å RMSD restraint.

Given the trimeric nature of MIF-2, three Ebselen molecules were docked at the corresponding sites in each monomer. The resulting complex was solvated in a TIP3P water box (55) with a 15 Å buffer, and Na+ and Cl− ions were added to neutralize the system at a physiological salt concentration of 0.15 M. MD simulations were performed using NAMD (56). To mimic the formation of a selenium–sulfur covalent bond, a harmonic restraint (2.23 Å, 388.24 kcal/mol/Å^2^) was applied between the selenium atom of Ebselen and the sulfur atom of Cys23. The system underwent a three-step equilibration: [1] relaxation of solvent only, [2] relaxation of solvent and side chains, and [3] relaxation of the entire system. Hydrogen mass repartitioning (HMR) (57) was applied to enable a 4 fs time step. Production MD simulations were run for 400 ns, while a parallel MD simulation of apo MIF-2 (no Ebselen bound) was performed under identical conditions for comparison. The X-ray maps were calculated from MD simulations using the sfall program in CCP4 (58). The apo and complex structures were then fitted to the maps to obtain the equilibrated structures. ProLif (59) was employed to analyze the protein-ligand interactions, and the results were visualized using PyMOL (60).

### Enzymatic Assays

MIF-2 activity was measured using a 100 mM stock solution of 4-hydroxyphenyl pyruvate (4-HPP) substrate, prepared in 500 mM ammonium acetate at pH 6.0 and rocked overnight to generate its keto form (61). MIF-2 enzymatic activity was determined by monitoring the increase in absorbance at 306 nm caused by enol–borate complex formation between boric acid and 4-HPP in the reaction solution. Absorbance was first recorded with a mixture of 1.2 mM 4-HPP and 420 mM boric acid, then the reaction was initiated by adding MIF-2 at a final concentration of 80 nM, and the absorbance was recorded during incubation for 3.5 min. To assess the inhibitory effect of Ebselen, the compound was first incubated with MIF-2 for one hour at room temperature. Absorbance of a “blank” sample was first recorded with a mixture of 1.2 mM 4-HPP and 420 mM boric acid and 160 nM Ebselen. The reaction was monitored by adding MIF-2 incubated with Ebselen to a fresh solution of 420 mM boric acid and 1.2 mM 4-HPP and the absorbance was recorded during incubation for 3.5 min. A control experiment was conducted using the same volume of DMSO used to solubilize Ebselen, but in the absence of the Ebselen molecule.

### Spin-down Assays

To quantify the percentage of MIF-2 precipitated from solution by Ebselen, 1 mL of 25 μM MIF-2 in a buffer of 20 mM Tris and 20 mM NaCl at pH 7.4 was centrifuged at 14,600 rpm for 5 minutes at room temperature. The soluble fraction was separated from any precipitate with a micropipette. Then, absorbance of the soluble fraction was measured at 280 nm using ε_280_ = 10500 M^−1^ cm^−1^, accounting for the molar extinction coefficient of Ebselen (30) to determine the amount of MIF-2 present at 0 hrs (initial reading), 6 hours, 12 hours, and 24 hours after addition of Ebselen at stoichiometric molar equivalence. A blank containing only buffer and the equivalent concentration of Ebselen was used for background correction. The assay was also carried out with 25 μM MIF-2 protein alone and MIF-2 protein with the same volume of DMSO used to solubilize Ebselen, which served as controls.

### Neutrophil Recruitment Assays

Wild-type adult male mice (10–12 weeks old) of genetic background strain C57BL6/J were purchased from Jackson Laboratory (Bar Harbor, ME) and housed in a pathogen-free animal facility at Cooper University Hospital, Camden, NJ, USA. Mice were administered a one-time intra-tracheal instillation of 50 μL of normal saline alone (vehicle) or 1 μg of either 1) Ebselen alone, 2) WT MIF2, 3) C56S MIF-2, or 4) a combination of MIF-2 and Ebselen, resuspended in 50 μl of DMSO and normal saline. Intra-tracheal instillation followed the methodology described prior work (13). The mice were sacrificed after 6 hours to collect bronchoalveolar lavage (BAL) and estimate the total protein content in the BAL fluid by BCA assay (Thermo Scientific, Rockford, IL). Approximately 1 mL of the BAL fluid was pelleted at 1000 rpm for 10 min at 4 °C. The pellet was resuspended in 200 μL of 1X PBS, and cytocentrifuged at 1000 rpm for 10 min at room temperature to spread the pellet evenly on slides as a smear. The slides were air dried and stained with Hema III differential Quick stain (Fisher Scientific, Cat. No. 122-911). The total percentage of neutrophils in the smears was determined manually following previous published methodology (21, 22). Briefly, following staining, the cytosmear was randomly divided into 4-5 areas that housed the maximum number of neutrophils. Two hundred different inflammatory cell types (including neutrophils) were counted in these areas, from which the percentage of neutrophils was calculated. At least 4-5 mice were used for the different groups of samples instilled. This animal study protocol was approved by the Institutional Animal Care and Use Committee of Cooper University Hospital, Camden, NJ, USA. Statistical analysis was done in GraphPad Prism 8.4.3 by one-way ANOVA with Tukey’s post-hoc correction, as appropriate.

## Acknowledgments

This work was supported by NIH grant R01 GM144451 (to GPL, VSB, and VB). VW was supported by NSF GRFP Grant 2040433.

## Data Availability

NMR assignments of MIF-2 have previously been deposited under BMRB entry 50790. All other data is available from the corresponding author upon request.

## Author Contributions

VW prepared MIF-2 samples performed NMR experiments, VW, SD, IV, and YL performed *in vitro* biochemical assays, SS and JW performed MD simulations, PD, XT, and DR performed *in vivo* neutrophil recruitment assays, VSB supervised computational work, VB supervised *in vivo* assays, and GPL conceived the study and supervised NMR studies. The manuscript was written through contributions of all authors.

## References

1. Merk M, Mitchell RA, Endres S, Bucala R. D-dopachrome tautomerase (D-DT or MIF-2): doubling the MIF cytokine family. Cytokine. 2012;59(1):10–7.

2. Chen E, Reiss K, Shah D, Manjula R, Allen B, Murphy EL, et al. A structurally preserved allosteric site in the MIF superfamily affects enzymatic activity and CD74 activation in D-dopachrome tautomerase. J Biol Chem. 2021;297(3):101061.

3. Chen E, Widjaja V, Kyro G, Allen B, Das P, Prahaladan VM, et al. Mapping N-to C-terminal allosteric coupling through disruption of a putative CD74 activation site in D-dopachrome tautomerase. J Biol Chem. 2023;299(6):104729.

4. Lubetsky JB, Dios A, Han J, Aljabari B, Ruzsicska B, Mitchell R, et al. The tautomerase active site of macrophage migration inhibitory factor is a potential target for discovery of novel anti-inflammatory agents. J Biol Chem. 2002;277(28):24976–82.

5. Trivedi-Parmar V, Jorgensen WL. Advances and Insights for Small Molecule Inhibition of Macrophage Migration Inhibitory Factor. J Med Chem. 2018;61(18):8104–19.

6. Winner M, Meier J, Zierow S, Rendon BE, Crichlow GV, Riggs R, et al. A novel, macrophage migration inhibitory factor suicide substrate inhibits motility and growth of lung cancer cells. Cancer Res. 2008;68(18):7253–7.

7. Cho Y, Crichlow GV, Vermeire JJ, Leng L, Du X, Hodsdon ME, et al. Allosteric inhibition of macrophage migration inhibitory factor revealed by ibudilast. Proc Natl Acad Sci U S A. 2010;107(25):11313–8.

8. Ouertatani-Sakouhi H, El-Turk F, Fauvet B, Cho MK, Pinar Karpinar D, Le Roy D, et al. Identification and characterization of novel classes of macrophage migration inhibitory factor (MIF) inhibitors with distinct mechanisms of action. J Biol Chem. 2010;285(34):26581–98.

9. Lange OF, Lakomek NA, Farès C, Schröder GF, Walter KF, Becker S, et al. Recognition dynamics up to microseconds revealed from an RDC-derived ubiquitin ensemble in solution. Science. 2008;320(5882):1471–5.

10. Shakhnovich E. Protein folding thermodynamics and dynamics: where physics, chemistry, and biology meet. Chem Rev. 2006;106(5):1559–88.

11. Tilstam PV, Schulte W, Holowka T, Kim BS, Nouws J, Sauler M, et al. MIF but not MIF-2 recruits inflammatory macrophages in an experimental polymicrobial sepsis model. J Clin Invest. 2021;131(23).

12. Fan C, Rajasekaran D, Syed MA, Leng L, Loria JP, Bhandari V, et al. MIF intersubunit disulfide mutant antagonist supports activation of CD74 by endogenous MIF trimer at physiologic concentrations. Proc Natl Acad Sci U S A. 2013;110(27):10994–9.

13. Parkins A, Chen E, Rangel VM, Singh M, Xue L, Lisi GP, et al. Ligand-induced conformational changes enable intersubunit communications in D-dopachrome tautomerase. Biophys J. 2023.

14. Lisi GP, Loria JP. Solution NMR Spectroscopy for the Study of Enzyme Allostery. Chem Rev. 2016;116(11):6323–69.

15. Wand AJ, Sharp KA. Measuring Entropy in Molecular Recognition by Proteins. Annu Rev Biophys. 2018;47:41–61.

16. Su XC, Jergic S, Ozawa K, Burns ND, Dixon NE, Otting G. Measurement of dissociation constants of high-molecular weight protein-protein complexes by transferred 15N-relaxation. J Biomol NMR. 2007;38(1):65–72.

17. Zhao Y, Hu J, Yang SS, Zhong J, Liu J, Wang S, et al. A redox switch regulates the assembly and anti-CRISPR activity of AcrIIC1. Nat Commun. 2022;13(1):7071.

18. Purslow JA, Khatiwada B, Bayro MJ, Venditti V. NMR Methods for Structural Characterization of Protein-Protein Complexes. Front Mol Biosci. 2020;7:9.

19. Cooke G, Armstrong ME, Donnelly SC. Macrophage migration inhibitory factor (MIF), enzymatic activity and the inflammatory response. Biofactors. 2009;35(2):165–8.

20. Parkins A, Skeens E, McCallum CM, Lisi GP, Pantouris G. The N-terminus of MIF regulates the dynamic profile of residues involved in CD74 activation. Biophys J. 2021;120(18):3893–900.

21. De Lorenzi D, Masserdotti C, Bertoncello D, Tranquillo V. Differential cell counts in canine cytocentrifuged bronchoalveolar lavage fluid: a study on reliable enumeration of each cell type. Vet Clin Pathol. 2009;38(4):532–6.

22. De Brauwer EI, Jacobs JA, Nieman F, Bruggeman CA, Drent M. Bronchoalveolar lavage fluid differential cell count. How many cells should be counted? Anal Quant Cytol Histol. 2002;24(6):337–41.

23. Parkins A, Das P, Prahaladan V, Rangel VM, Xue L, Sankaran B, et al. 2,5-Pyridinedicarboxylic acid is a bioactive and highly selective inhibitor of D-dopachrome tautomerase. Structure. 2023;31(3):355–67.e4.

24. Schinagl A, Kerschbaumer RJ, Sabarth N, Douillard P, Scholz P, Voelkel D, et al. Role of the Cysteine 81 Residue of Macrophage Migration Inhibitory Factor as a Molecular Redox Switch. Biochemistry. 2018;57(9):1523–32.

25. Skeens E, Pantouris G, Shah D, Manjula R, Ombrello MJ, Maluf NK, et al. A Cysteine Variant at an Allosteric Site Alters MIF Dynamics and Biological Function in Homo- and Heterotrimeric Assemblies. Front Mol Biosci. 2022;9:783669.

26. Skeens E, Gadzuk-Shea M, Shah D, Bhandari V, Schweppe DK, Berlow RB, et al. Redox-dependent structure and dynamics of macrophage migration inhibitory factor reveal sites of latent allostery. Structure. 2022;30(6):840–50.e6.

27. Coleman AM, Rendon BE, Zhao M, Qian MW, Bucala R, Xin D, et al. Cooperative regulation of non-small cell lung carcinoma angiogenic potential by macrophage migration inhibitory factor and its homolog, D-dopachrome tautomerase. J Immunol. 2008;181(4):2330–7.

28. Feng Q, Li X, Sun W, Li Y, Yuan Y, Guan B, et al. Discovery of Ebselen as an Inhibitor of 6PGD for Suppressing Tumor Growth. Cancer Manag Res. 2020;12:6921–34.

29. Landgraf AD, Alsegiani AS, Alaqel S, Thanna S, Shah ZA, Sucheck SJ. Neuroprotective and Anti-neuroinflammatory Properties of Ebselen Derivatives and Their Potential to Inhibit Neurodegeneration. ACS Chem Neurosci. 2020;11(19):3008–16.

30. Zhao R, Holmgren A. A novel antioxidant mechanism of ebselen involving ebselen diselenide, a substrate of mammalian thioredoxin and thioredoxin reductase. J Biol Chem. 2002;277(42):39456–62.

31. Zhao R, Masayasu H, Holmgren A. Ebselen: a substrate for human thioredoxin reductase strongly stimulating its hydroperoxide reductase activity and a superfast thioredoxin oxidant. Proc Natl Acad Sci U S A. 2002;99(13):8579–84.

32. Capper MJ, Wright GSA, Barbieri L, Luchinat E, Mercatelli E, McAlary L, et al. The cysteine-reactive small molecule ebselen facilitates effective SOD1 maturation. Nat Commun. 2018;9(1):1693.

33. Pantouris G, Syed MA, Fan C, Rajasekaran D, Cho TY, Rosenberg EM, Jr., et al. An Analysis of MIF Structural Features that Control Functional Activation of CD74. Chem Biol. 2015;22(9):1197–205.

34. Pantouris G, Bucala R, Lolis EJ. Structural Plasticity in the C-Terminal Region of Macrophage Migration Inhibitory Factor-2 Is Associated with an Induced Fit Mechanism for a Selective Inhibitor. Biochemistry. 2018;57(26):3599–605.

35. Schinagl A, Thiele M, Douillard P, Völkel D, Kenner L, Kazemi Z, et al. Oxidized macrophage migration inhibitory factor is a potential new tissue marker and drug target in cancer. Oncotarget. 2016;7(45):73486–96.

36. Thiele M, Bernhagen J. Link between macrophage migration inhibitory factor and cellular redox regulation. Antioxid Redox Signal. 2005;7(9-10):1234–48.

37. Sajko S, Skeens E, Schinagl A, Ferhat M, Mirkina I, Mayer J, et al. Redox-dependent plasticity of oxMIF facilitates its interaction with CD74 and therapeutic antibodies. Redox Biol. 2024;75:103264.

38. Tilstam PV, Pantouris G, Corman M, Andreoli M, Mahboubi K, Davis G, et al. A selective small-molecule inhibitor of macrophage migration inhibitory factor-2 (MIF-2), a MIF cytokine superfamily member, inhibits MIF-2 biological activity. J Biol Chem. 2019;294(49):18522–31.

39. Al-Abed Y, Dabideen D, Aljabari B, Valster A, Messmer D, Ochani M, et al. ISO-1 binding to the tautomerase active site of MIF inhibits its pro-inflammatory activity and increases survival in severe sepsis. J Biol Chem. 2005;280(44):36541–4.

40. Xiao Z, Osipyan A, Song S, Chen D, Schut RA, van Merkerk R, et al. Thieno[2,3-d]pyrimidine-2,4(1H,3H)-dione Derivative Inhibits d-Dopachrome Tautomerase Activity and Suppresses the Proliferation of Non-Small Cell Lung Cancer Cells. J Med Chem. 2022;65(3):2059–77.

41. Shi X, Leng L, Wang T, Wang W, Du X, Li J, et al. CD44 is the signaling component of the macrophage migration inhibitory factor-CD74 receptor complex. Immunity. 2006;25(4):595–606.

42. Delaglio F, Grzesiek S, Vuister GW, Zhu G, Pfeifer J, Bax A. NMRPipe: a multidimensional spectral processing system based on UNIX pipes. J Biomol NMR. 1995;6(3):277–93.

43. Lee W, Tonelli M, Markley JL. NMRFAM-SPARKY: enhanced software for biomolecular NMR spectroscopy. Bioinformatics. 2015;31(8):1325–7.

44. Grzesiek S, Stahl SJ, Wingfield PT, Bax A. The CD4 determinant for downregulation by HIV-1 Nef directly binds to Nef. Mapping of the Nef binding surface by NMR. Biochemistry. 1996;35(32):10256–61.

45. Zhu G, Xia Y, Nicholson LK, Sze KH. Protein dynamics measurements by TROSY-based NMR experiments. J Magn Reson. 2000;143(2):423–6.

46. Kay LE, Torchia DA, Bax A. Backbone dynamics of proteins as studied by 15N inverse detected heteronuclear NMR spectroscopy: application to staphylococcal nuclease. Biochemistry. 1989;28(23):8972–9.

47. Trott O, Olson AJ. AutoDock Vina: improving the speed and accuracy of docking with a new scoring function, efficient optimization, and multithreading. J Comput Chem. 2010;31(2):455–61.

48. Pettersen EF, Goddard TD, Huang CC, Couch GS, Greenblatt DM, Meng EC, et al. UCSF Chimera--a visualization system for exploratory research and analysis. J Comput Chem. 2004;25(13):1605–12.

49. Malde AK, Zuo L, Breeze M, Stroet M, Poger D, Nair PC, et al. An Automated Force Field Topology Builder (ATB) and Repository: Version 1.0. J Chem Theory Comput. 2011;7(12):4026–37.

50. Case DA, Aktulga HM, Belfon K, Cerutti DS, Cisneros GA, Cruzeiro VWD, et al. AmberTools. Journal of Chemical Information and Modeling. 2023;63(20):6183–91.

51. Case DA, Aktulga, H. M., Belfon, K., Ben-Shalom, I. Y., Berryman, J. T., Brozell, S. R., Cerutti, D. S., Cheatham, T. E., III, Cisneros, G. A., Cruzeiro, V. W. D., Darden, T. A., Forouzesh, N., Ghazimirsaeed, M., Giambaşu, G., Giese, T., Gilson, M. K., Gohlke, H., Goetz, A. W., Harris, J., Huang, Z., Izadi, S., Izmailov, S. A., Kasavajhala, K., Kaymak, M. C., Kolossváry, I., Kovalenko, A., Kurtzman, T., Lee, T. S., Li, P., Li, Z., Lin, C., Liu, J., Luchko, T., Luo, R., Machado, M., Manathunga, M. K., Merz, K. M., Miao, Y., Mikhailovskii, O., Monard, G., Nguyen, H., O’Hearn, K. A., Onufriev, A., Pan, F., Pantano, S., Rahnamoun, A., Roe, D. R., Roitberg, A., Sagui, C., Schott-Verdugo, S., Shajan, A., Shen, J., Simmerling, C. L., Skrynnikov, N. R., Smith, J., Swails, J., Walker, R. C., Wang, J., Wang, X., Wu, Y., Xiong, Y., Xue, Y., York, D. M., Zhao, C., Zhu, Q., & Kollman, P. A. AMBER. University of California, San Francisco; 2025.

52. 2025-2 SR. New York, NY: Maestro. Schrödinger, LLC.

53. Olsson MH, Søndergaard CR, Rostkowski M, Jensen JH. PROPKA3: Consistent Treatment of Internal and Surface Residues in Empirical pKa Predictions. J Chem Theory Comput. 2011;7(2):525–37.

54. Jorgensen WL, Tirado-Rives J. The OPLS [optimized potentials for liquid simulations] potential functions for proteins, energy minimizations for crystals of cyclic peptides and crambin. J Am Chem Soc. 1988;110(6):1657–66.

55. Klein WLJJCJDMRWIML. Comparison of simple potential functions for simulating liquid wate. J Chem Phys. 1983;79:926–36.

56. Phillips JC, Braun R, Wang W, Gumbart J, Tajkhorshid E, Villa E, et al. Scalable molecular dynamics with NAMD. J Comput Chem. 2005;26(16):1781–802.

57. Hopkins CW, Le Grand S, Walker RC, Roitberg AE. Long-Time-Step Molecular Dynamics through Hydrogen Mass Repartitioning. J Chem Theory Comput. 2015;11(4):1864–74.

58. Winn MD, Ballard CC, Cowtan KD, Dodson EJ, Emsley P, Evans PR, et al. Overview of the CCP4 suite and current developments. Acta Crystallogr D Biol Crystallogr. 2011;67(Pt 4):235–42.

59. Schneider G, et al. ProLif: An interactive software tool for protein-ligand interaction analysis. J Comput Aided Mol Des. 2009;23(2):121–32.

60. DeLano WL. PyMOL: An open-source molecular graphics tool. CCP4 Newsletter on Protein Crystallography. 2002;40:82–9.

61. Leng L, Metz CN, Fang Y, Xu J, Donnelly S, Baugh J, et al. MIF signal transduction initiated by binding to CD74. J Exp Med. 2003;197(11):1467–76.

