## Supporting Information for "Allosteric Modulation of MIF-2 Structure, Catalysis, and Biological Signaling via Cysteine Residues and a Small Molecule, Ebselen"

### List of Tables and Figures

**Figure S1.** Effect of cysteine mutations on MIF-2 structure monitored by NMR
**Figure S2.** Circular dichroism and thermal stability of WT MIF-2 and variants
**Figure S3.** Effect of Ebselen on MIF-2 and variants monitored by NMR
**Figure S4.** Reduction of MIF-2 cysteines attenuates the structural impact of Ebselen
**Figure S5.** Ebelsen-induced RMSF from MD simulations
**Figure S6.** $R_1$, $R_2$, and $R_2/R_1$ values for WT MIF-2 (±Ebselen) and C56S MIF-2 (±Ebselen)
**Figure S7.** Cytosmears of neutrophils from CD74-mediated inflammatory activity of MIF-2
**Table S1.** $R_1$, $R_2$, and $R_2/R_1$ values for WT MIF-2
**Table S2.** $R_1$, $R_2$, and $R_2/R_1$ values for WT MIF-2 bound to Ebselen
**Table S3.** $R_1$, $R_2$, and $R_2/R_1$ values for C56S MIF-2
**Table S4.** $R_1$, $R_2$, and $R_2/R_1$ values for C56S MIF-2 bound to Ebselen

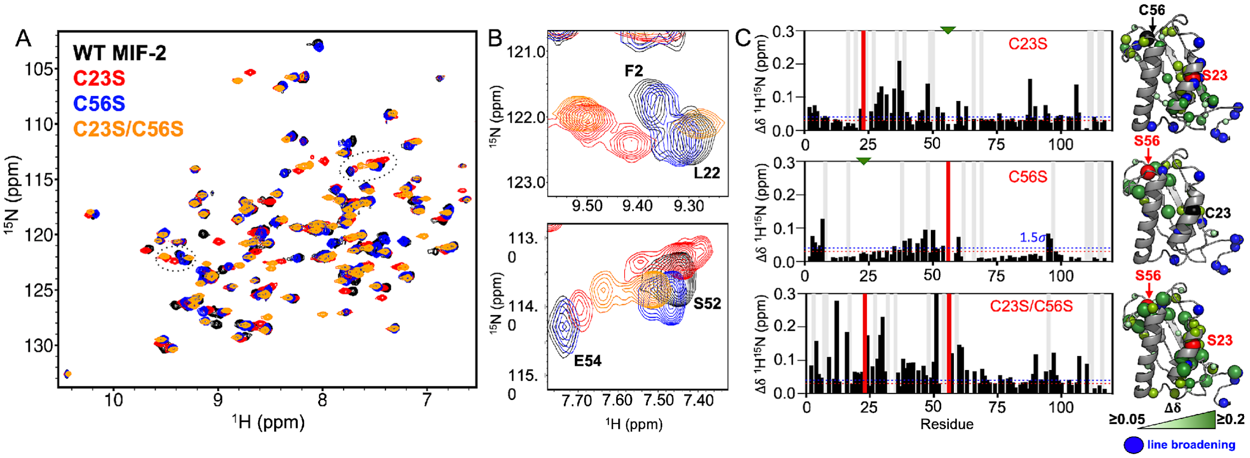


**Figure S1.** Effect of cysteine mutations on MIF-2 structure. **(A)** $^{1}H$-$^{15}N$ HSQC NMR spectra of WT MIF-2 (black), C23S MIF-2 (red), C56S MIF-2 (blue), and C23S/C56S MIF-2 (orange). Representative spectral regions highlighted by dashed circles are shown in close view in **(B)**, where local fluctuations in atomic environment are apparent. **(C)** Per-residue NMR CSPs (black bars) caused by MIF-2 mutations, relative to WT MIF-2. Light gray bars denote sites of NMR line broadening. Red bars denote a Cys-to-Ser mutation, and green triangles denote a native Cys residue. CSPs >1.5σ of the 10% trimmed mean of all shifts are mapped on the MIF-2 monomer (green spheres). Sphere size correlates with the intensity of the CSP. Blue spheres represent sites of NMR line broadening, red spheres denote a Cys-to-Ser mutation, and black spheres denote a native Cys residue.

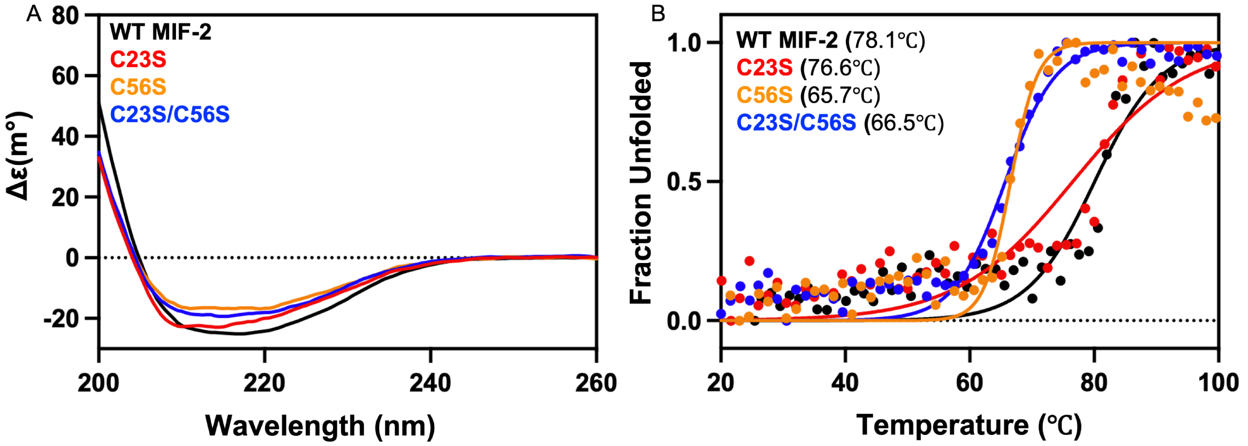


**Figure S2.** Circular dichroism assessment of MIF-2 secondary structure and thermal stability of WT MIF-2 and mutants. **(A)** Far-UV circular dichroism spectra suggesting a mix of α-β structure expected for MIF-2, which is retained upon mutation of cysteine residues. **(B)** Thermal unfolding of WT MIF-2 and variants (λ = 218 nm) revealing that C23S (red) is similarly thermostable, while C56S (orange) and C23S/C56S (blue) are substantially destabilized.

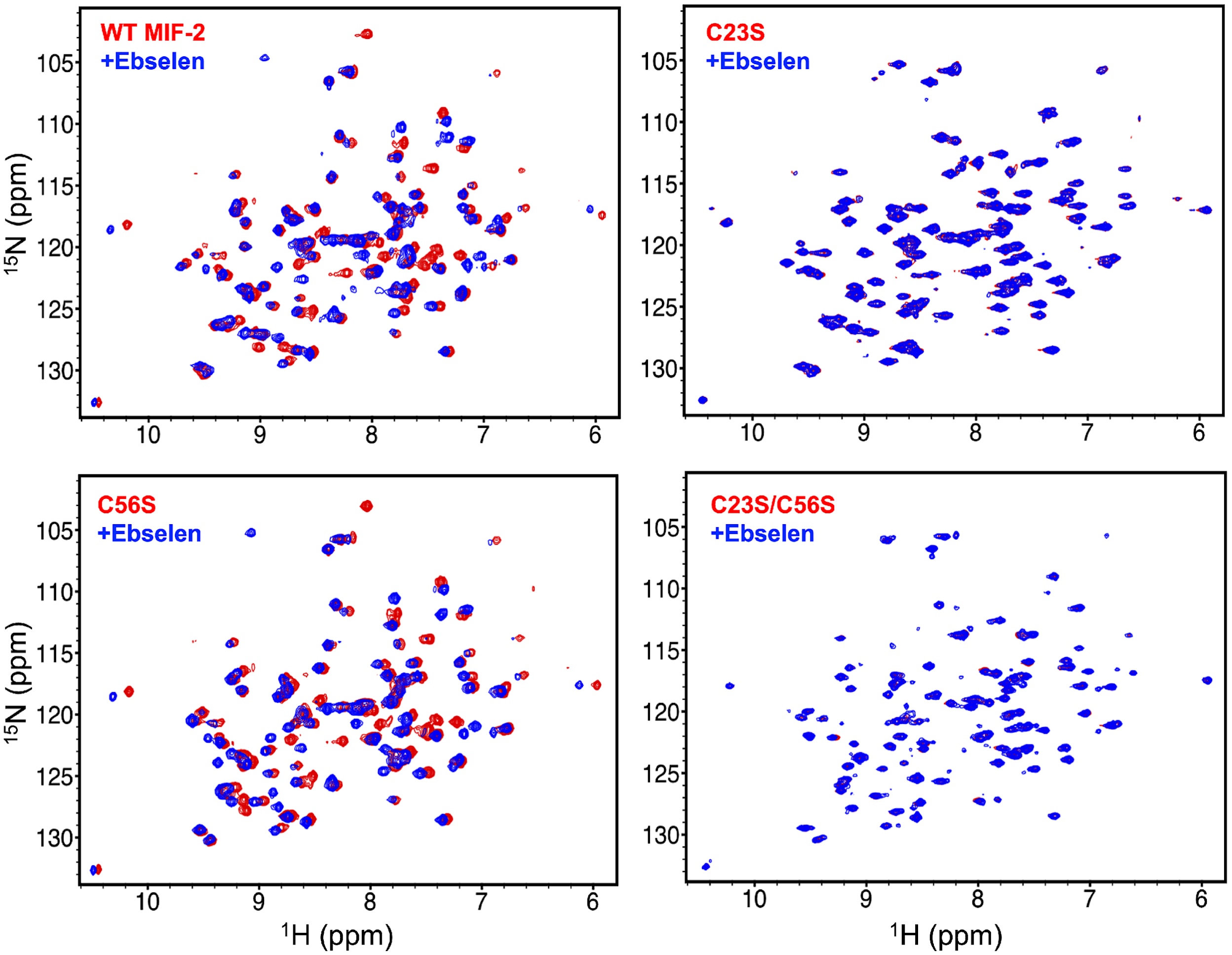


**Figure S3.** $^{1}H$-$^{15}N$ HSQC NMR spectra of WT-MIF-2 and MIF-2 variants in the absence (red) and presence (blue) of Ebselen. Data were collected at 14.1 T and 303 K using 0.5 mM MIF-2 in a buffer of 20 mM Tris, 20 mM NaCl, 1.5% (v/v) DMSO-d6, and 10% $D_2O$ at pH 7.4.

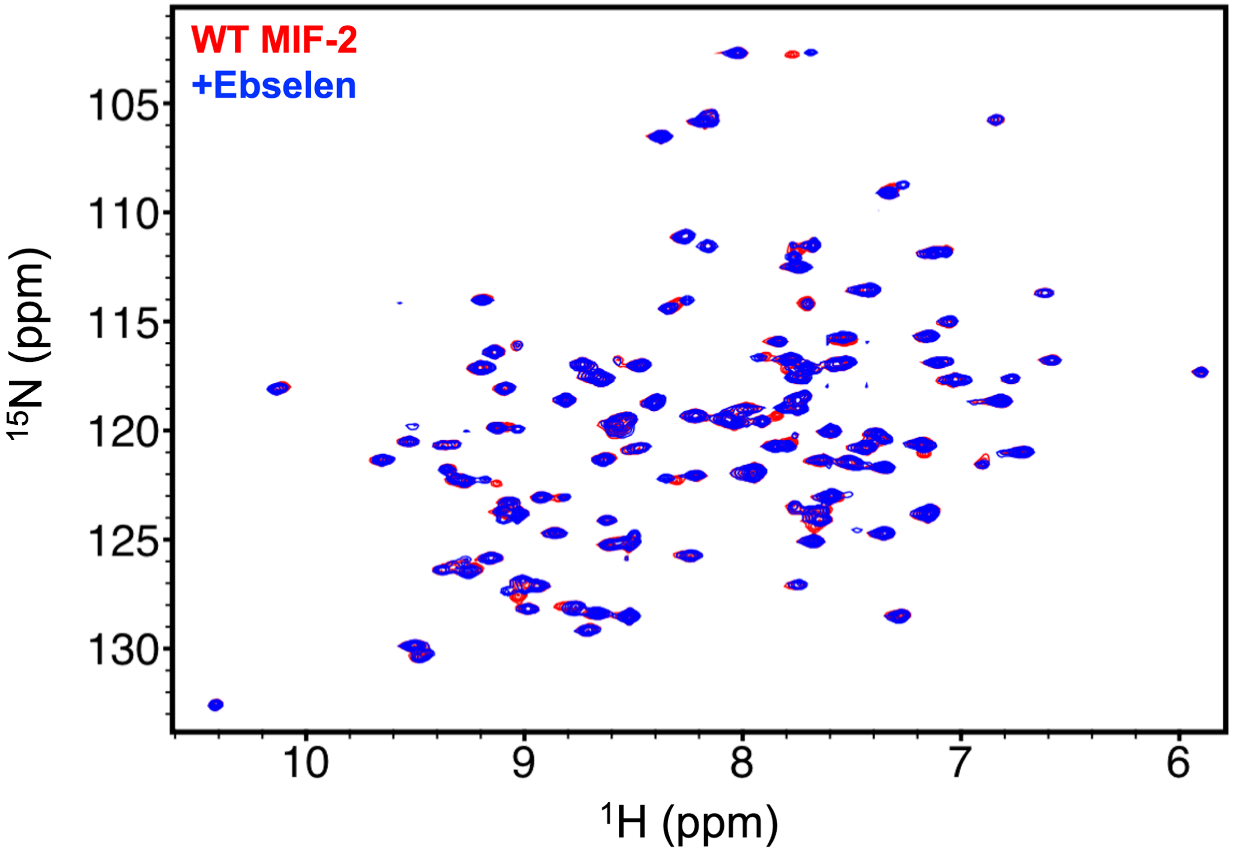


**Figure S4.** $^{1}H$-$^{15}N$ HSQC NMR spectra of WT-MIF-2 in the absence (red) and presence (blue) of Ebselen in the presence of 5 mM DTT. Data were collected at 14.1 T and 303 K using 0.5 mM MIF-2 in a buffer of 20 mM Tris, 20 mM NaCl, 1.5% (v/v) DMSO-d6, and 10% $D_2O$ at pH 7.4.

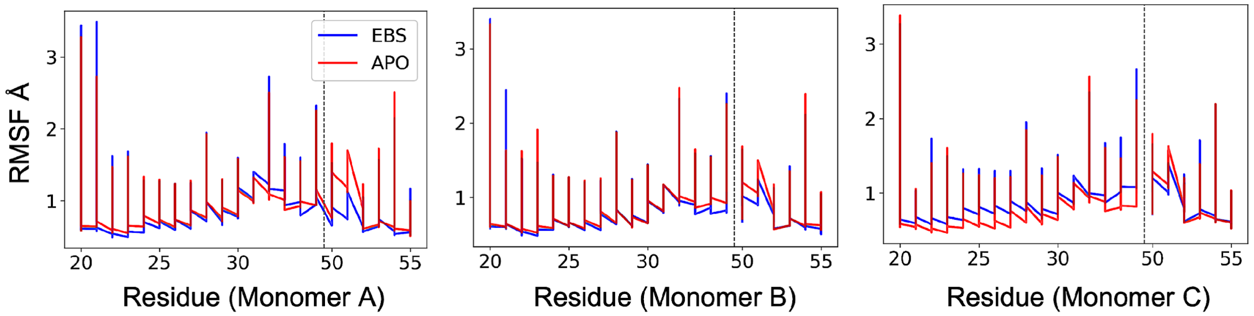


**Figure S5.** Ebelsen-induced root-mean-square fluctuations (RMSF, Å) of residues surrounding the Ebselen binding sites in each of the MIF-2 monomers. Average RMSF values from the entire MD trajectories are reported for apo MIF-2 (red) and Ebselen-bound MIF-2 (blue).

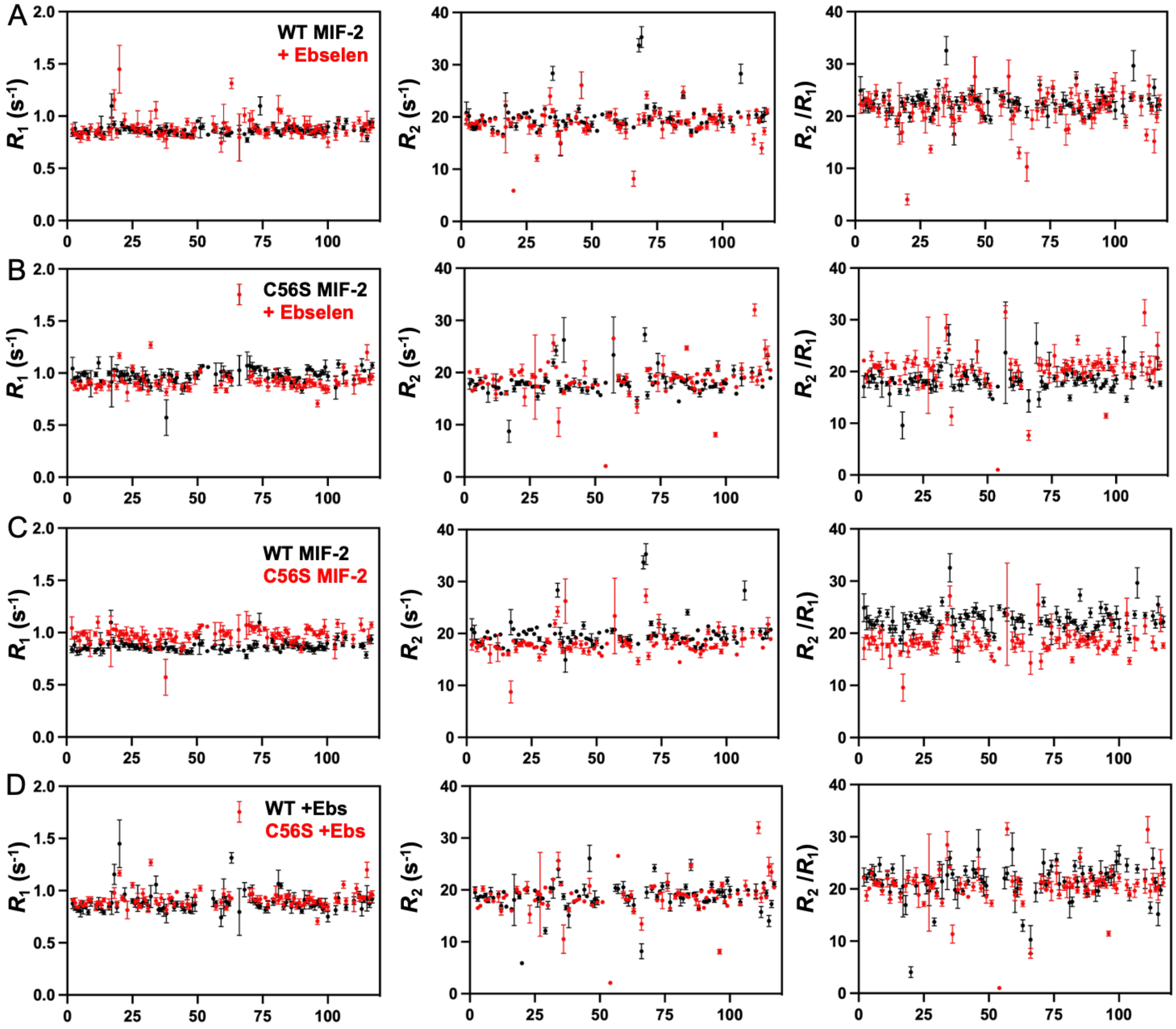


**Figure S6.** NMR spin relaxation parameters for WT and C56S MIF-2. **(A)** Per-residue $R_1$, $R_2$, and $R_2/R_1$ values measured at 14.1 T (600 MHz Larmor frequency) for WT MIF-2 (black) and WT MIF-2 bound to Ebselen (red). **(B)** The same parameters measured for C56S MIF-2 (black) and C56S MIF-2 bound to Ebselen (red). **(C)** Comparison of $R_1$, $R_2$, and $R_2/R_1$ values for WT MIF-2 (black) and C56S MIF-2 (red). **(D)** Comparison of the same parameters for each protein bound to Ebselen.

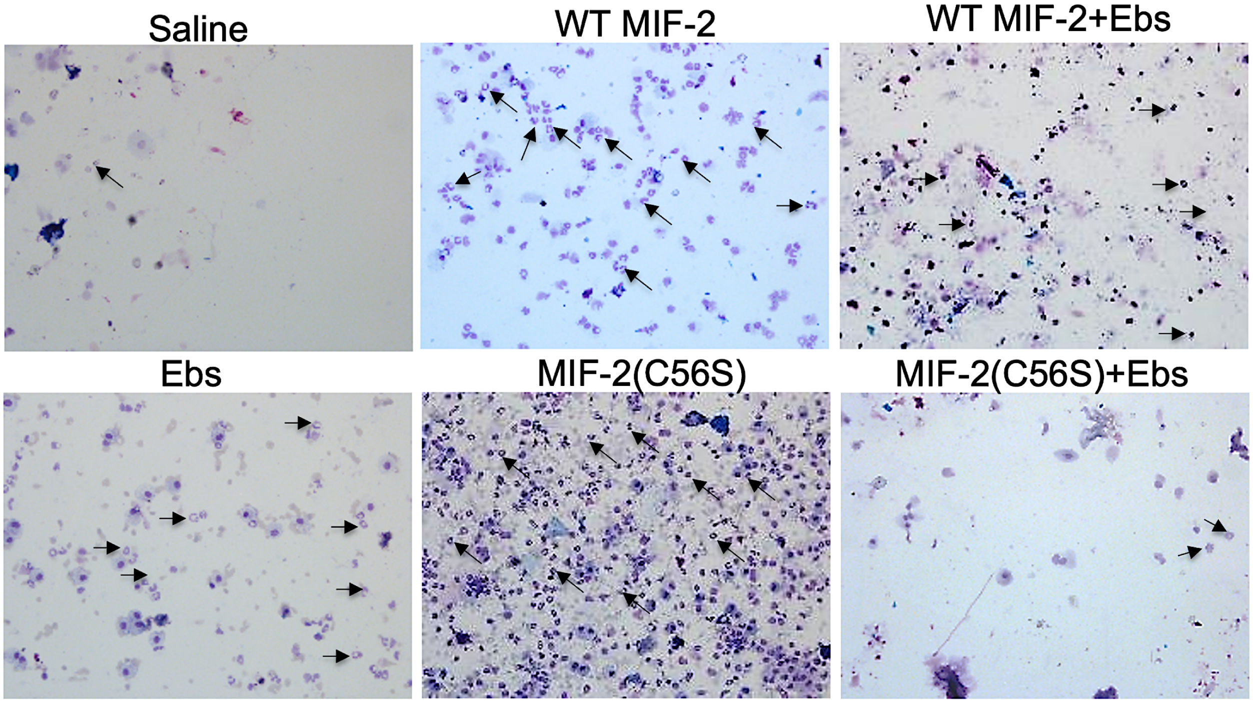


**Figure S7.** Ebselen modulates CD74-mediated inflammatory activity of WT- and C56S MIF-2 *in vivo*. Bronchoalveolar lavage (BAL) cytosmears showing neutrophil recruitment (arrows) by WT- and C56S MIF-2 in the absence and presence of Ebselen. Saline and Ebselen alone are shown as controls.

**Table S1.** $R_1$, $R_2$, and $R_2/R_1$ values for WT MIF-2

| Residue | *R1* | | *R2* | | *R2/R1* | | Residue | *R1* | | *R2* | | *R2/R1* | |
|---|---|---|---|---|---|---|---|---|---|---|---|---|---|
| # | Mean | Error | Mean | Error | Mean | Error | # | Mean | Error | Mean | Error | Mean | Error |
| 2 | 0.838 | 0.030 | 20.833 | 2.036 | 24.875 | 2.689 | 57 | 0.880 | 0.044 | 20.743 | 0.362 | 23.584 | 1.264 |
| 3 | 0.880 | 0.061 | 20.190 | 1.410 | 22.936 | 2.281 | 59 | 0.838 | 0.047 | 19.186 | 0.456 | 22.909 | 1.355 |
| 4 | 0.820 | 0.038 | 18.228 | 0.704 | 22.220 | 1.252 | 60 | 0.880 | 0.055 | 18.440 | 0.972 | 20.948 | 1.685 |
| 5 | 0.825 | 0.034 | 17.867 | 0.517 | 21.654 | 1.021 | 61 | 0.820 | 0.012 | 18.539 | 0.344 | 22.599 | 0.504 |
| 6 | 0.842 | 0.011 | 18.159 | 0.534 | 21.573 | 0.642 | 62 | 0.825 | 0.022 | 18.868 | 0.566 | 22.868 | 0.864 |
| 7 | 0.861 | 0.023 | 19.260 | 0.275 | 22.381 | 0.661 | 63 | 0.842 | 0.017 | 18.723 | 1.052 | 22.243 | 1.256 |
| 8 | 0.845 | 0.021 | 20.321 | 0.374 | 24.060 | 0.742 | 66 | 0.867 | 0.044 | 18.015 | 0.266 | 20.771 | 1.078 |
| 9 | 0.887 | 0.037 | 19.044 | 0.131 | 21.463 | 0.914 | 68 | 0.828 | 0.013 | 33.727 | 1.285 | 40.742 | 2.704 |
| 11 | 0.824 | 0.026 | 19.612 | 0.143 | 23.789 | 0.726 | 69 | 0.773 | 0.022 | 35.298 | 2.031 | 45.641 | 4.799 |
| 12 | 0.843 | 0.014 | 17.541 | 0.646 | 20.803 | 0.749 | 70 | 0.861 | 0.021 | 19.216 | 0.447 | 22.329 | 0.728 |
| 13 | 0.889 | 0.110 | 19.802 | 0.194 | 22.277 | 2.825 | 71 | 0.845 | 0.024 | 21.959 | 0.440 | 25.999 | 0.911 |
| 14 | 0.826 | 0.029 | 17.182 | 0.133 | 20.808 | 0.703 | 72 | 0.887 | 0.039 | 19.857 | 0.438 | 22.379 | 1.116 |
| 16 | 0.888 | 0.036 | 16.639 | 0.155 | 18.735 | 0.784 | 73 | 0.888 | 0.031 | 19.531 | 0.108 | 21.992 | 0.784 |
| 17 | 1.097 | 0.118 | 22.188 | 2.432 | 20.217 | 3.699 | 74 | 1.097 | 0.087 | 21.925 | 0.769 | 19.978 | 2.154 |
| 18 | 0.921 | 0.032 | 20.864 | 0.250 | 22.658 | 0.892 | 76 | 0.869 | 0.016 | 20.747 | 1.033 | 23.880 | 1.317 |
| 19 | 0.864 | 0.017 | 18.443 | 0.463 | 21.357 | 0.648 | 77 | 0.849 | 0.028 | 18.957 | 0.385 | 22.332 | 0.832 |
| 20 | 0.847 | 0.022 | 19.790 | 0.374 | 23.372 | 0.736 | 79 | 0.921 | 0.022 | 19.391 | 0.530 | 21.059 | 0.781 |
| 21 | 0.917 | 0.028 | 18.172 | 0.717 | 19.807 | 0.954 | 80 | 0.864 | 0.029 | 19.912 | 1.142 | 23.059 | 1.526 |
| 22 | 0.882 | 0.023 | 20.362 | 0.316 | 23.091 | 0.708 | 81 | 0.847 | 0.021 | 19.440 | 0.775 | 22.959 | 1.051 |
| 23 | 0.894 | 0.014 | 19.916 | 0.341 | 22.286 | 0.526 | 82 | 0.815 | 0.011 | 18.086 | 0.226 | 22.192 | 0.379 |
| 24 | 0.869 | 0.034 | 20.721 | 0.167 | 23.850 | 0.958 | 83 | 0.862 | 0.020 | 19.109 | 0.110 | 22.167 | 0.526 |
| 25 | 0.889 | 0.031 | 20.760 | 0.309 | 23.355 | 0.916 | 84 | 0.917 | 0.023 | 19.124 | 0.240 | 20.845 | 0.617 |
| 26 | 0.862 | 0.029 | 19.924 | 0.263 | 23.112 | 0.828 | 85 | 0.882 | 0.029 | 24.091 | 0.529 | 27.319 | 1.160 |
| 27 | 0.867 | 0.025 | 21.146 | 0.271 | 24.403 | 0.774 | 86 | 0.894 | 0.021 | 18.811 | 0.222 | 21.050 | 0.552 |
| 28 | 0.842 | 0.013 | 18.936 | 0.441 | 22.496 | 0.597 | 87 | 0.869 | 0.023 | 18.961 | 0.281 | 21.824 | 0.662 |
| 29 | 0.877 | 0.020 | 17.010 | 0.324 | 19.391 | 0.546 | 88 | 0.889 | 0.020 | 17.268 | 0.423 | 19.427 | 0.601 |
| 30 | 0.858 | 0.035 | 17.581 | 0.760 | 20.499 | 1.143 | 89 | 0.862 | 0.032 | 20.610 | 0.291 | 23.908 | 0.950 |
| 31 | 0.846 | 0.026 | 18.365 | 0.310 | 21.708 | 0.740 | 90 | 0.867 | 0.015 | 17.265 | 0.358 | 19.924 | 0.499 |
| 32 | 0.828 | 0.023 | 21.552 | 0.382 | 26.034 | 0.847 | 91 | 0.842 | 0.019 | 17.771 | 0.309 | 21.112 | 0.566 |
| 34 | 0.848 | 0.016 | 19.956 | 0.266 | 23.528 | 0.531 | 92 | 0.877 | 0.034 | 18.829 | 0.251 | 21.465 | 0.892 |
| 35 | 0.871 | 0.042 | 28.361 | 1.351 | 32.558 | 2.710 | 93 | 0.858 | 0.040 | 21.368 | 0.267 | 24.915 | 1.185 |
| 36 | 0.865 | 0.029 | 20.032 | 1.192 | 23.157 | 1.590 | 94 | 0.846 | 0.032 | 18.968 | 0.165 | 22.420 | 0.847 |
| 37 | 0.850 | 0.019 | 18.342 | 1.083 | 21.588 | 1.262 | 95 | 0.828 | 0.032 | 20.408 | 0.118 | 24.653 | 0.912 |
| 38 | 0.898 | 0.037 | 14.914 | 2.380 | 16.600 | 2.106 | 96 | 0.848 | 0.017 | 19.829 | 0.297 | 23.379 | 0.584 |
| 39 | 0.840 | 0.030 | 19.212 | 0.151 | 22.882 | 0.804 | 97 | 0.871 | 0.041 | 19.658 | 0.359 | 22.567 | 1.147 |
| 40 | 0.859 | 0.015 | 18.501 | 0.479 | 21.536 | 0.638 | 98 | 0.865 | 0.033 | 18.904 | 0.230 | 21.853 | 0.875 |
| 41 | 0.850 | 0.025 | 20.080 | 0.480 | 23.635 | 0.885 | 99 | 0.850 | 0.024 | 20.313 | 0.953 | 23.908 | 1.321 |
| 42 | 0.826 | 0.022 | 19.172 | 0.216 | 23.217 | 0.637 | 100 | 0.898 | 0.054 | 18.975 | 0.983 | 21.120 | 1.682 |
| 44 | 0.824 | 0.033 | 19.822 | 0.253 | 24.044 | 0.966 | 103 | 0.885 | 0.049 | 19.406 | 0.674 | 21.929 | 1.435 |
| 45 | 0.869 | 0.019 | 21.556 | 0.711 | 24.811 | 1.046 | 104 | 0.960 | 0.036 | 18.225 | 0.212 | 18.990 | 0.807 |
| 46 | 0.815 | 0.024 | 19.091 | 0.234 | 23.425 | 0.700 | 106 | 0.887 | 0.020 | 20.929 | 0.393 | 23.587 | 0.724 |
| 47 | 0.815 | 0.013 | 18.907 | 0.345 | 23.199 | 0.527 | 107 | 0.953 | 0.027 | 28.281 | 1.848 | 29.666 | 2.901 |
| 48 | 0.822 | 0.030 | 18.646 | 1.314 | 22.674 | 1.692 | 110 | 0.866 | 0.015 | 20.263 | 0.328 | 23.404 | 0.559 |
| 49 | 0.885 | 0.019 | 17.775 | 0.404 | 20.085 | 0.592 | 113 | 0.875 | 0.021 | 20.408 | 0.658 | 23.327 | 0.962 |
| 50 | 0.953 | 0.011 | 18.608 | 0.582 | 19.520 | 0.621 | 114 | 0.874 | 0.030 | 19.066 | 0.333 | 21.811 | 0.829 |
| 51 | 0.889 | 0.097 | 20.149 | 0.318 | 22.668 | 2.562 | 115 | 0.785 | 0.031 | 19.992 | 1.039 | 25.470 | 1.607 |
| 52 | 0.901 | 0.030 | 17.358 | 0.253 | 19.267 | 0.711 | 116 | 0.933 | 0.026 | 20.450 | 1.330 | 21.922 | 1.604 |
| 54 | 0.822 | 0.018 | 20.492 | 0.142 | 24.918 | 0.544 | 117 | 0.935 | 0.042 | 20.747 | 0.215 | 22.178 | 1.088 |
| 56 | 0.857 | 0.013 | 20.500 | 0.551 | 23.924 | 0.751 | | | | | | | |

**Table S2.** $R_1$, $R_2$, and $R_2/R_1$ values for WT MIF-2 bound to Ebselen

| Residue | *R1* | | *R2* | | *R2/R1* | |
|---|---|---|---|---|---|---|
| # | Mean | Error | Mean | Error | Mean | Error |
| 2 | 0.876 | 0.048 | 19.531 | 1.076 | 22.285 | 1.735 |
| 3 | 0.831 | 0.020 | 18.328 | 0.514 | 22.049 | 0.778 |
| 4 | 0.816 | 0.024 | 18.734 | 0.244 | 22.967 | 0.682 |
| 5 | 0.804 | 0.055 | 18.671 | 0.359 | 23.226 | 1.485 |
| 6 | 0.839 | 0.064 | 18.993 | 0.426 | 22.640 | 1.704 |
| 7 | 0.839 | 0.012 | 17.953 | 0.393 | 21.400 | 0.530 |
| 8 | 0.792 | 0.026 | 19.569 | 0.923 | 24.697 | 1.386 |
| 9 | 0.888 | 0.037 | 18.471 | 0.392 | 20.798 | 0.966 |
| 11 | 0.848 | 0.026 | 19.220 | 0.189 | 22.660 | 0.705 |
| 12 | 0.874 | 0.060 | 16.935 | 0.660 | 19.373 | 1.468 |
| 13 | 0.822 | 0.033 | 19.619 | 0.236 | 23.877 | 0.920 |
| 14 | 0.793 | 0.023 | 16.787 | 0.161 | 21.168 | 0.570 |
| 16 | 0.848 | 0.033 | 16.321 | 0.354 | 19.243 | 0.797 |
| 17 | 0.882 | 0.110 | 18.090 | 4.941 | 20.514 | 5.878 |
| 18 | 1.155 | 0.097 | 19.543 | 0.190 | 16.926 | 1.834 |
| 19 | 0.839 | 0.047 | 18.720 | 0.442 | 22.314 | 1.268 |
| 20 | 1.450 | 0.229 | 5.882 | 0.156 | 4.057 | 1.038 |
| 22 | 0.964 | 0.064 | 19.877 | 0.597 | 20.612 | 1.603 |
| 25 | 0.900 | 0.023 | 19.885 | 0.379 | 22.092 | 0.722 |
| 26 | 0.874 | 0.032 | 18.238 | 1.533 | 20.864 | 1.829 |
| 27 | 0.925 | 0.125 | 18.142 | 0.533 | 19.612 | 2.789 |
| 28 | 0.857 | 0.025 | 18.365 | 0.128 | 21.433 | 0.619 |
| 29 | 0.886 | 0.041 | 12.108 | 0.613 | 13.670 | 0.759 |
| 30 | 0.881 | 0.013 | 17.044 | 0.750 | 19.345 | 0.808 |
| 31 | 0.858 | 0.054 | 20.371 | 0.333 | 23.752 | 1.501 |
| 32 | 0.945 | 0.105 | 20.272 | 1.886 | 21.447 | 3.278 |
| 34 | 1.058 | 0.083 | 23.987 | 1.588 | 22.679 | 2.819 |
| 35 | 0.818 | 0.042 | 21.204 | 0.404 | 25.912 | 1.320 |
| 36 | 0.818 | 0.020 | 18.713 | 0.571 | 22.867 | 0.856 |
| 37 | 0.836 | 0.082 | 16.573 | 0.431 | 19.821 | 1.879 |
| 38 | 0.781 | 0.088 | 15.092 | 2.369 | 19.333 | 3.063 |
| 40 | 0.904 | 0.026 | 17.690 | 0.124 | 19.565 | 0.576 |
| 41 | 0.838 | 0.032 | 19.712 | 0.256 | 23.517 | 0.886 |
| 42 | 0.871 | 0.026 | 18.737 | 0.358 | 21.510 | 0.742 |
| 44 | 0.845 | 0.038 | 20.036 | 1.084 | 23.723 | 1.674 |
| 45 | 0.815 | 0.029 | 18.786 | 0.162 | 23.051 | 0.780 |
| 46 | 0.946 | 0.034 | 26.069 | 2.535 | 27.555 | 3.807 |
| 47 | 0.797 | 0.018 | 17.437 | 0.468 | 21.883 | 0.692 |
| 48 | 0.858 | 0.079 | 18.232 | 1.622 | 21.258 | 2.610 |
| 49 | 0.867 | 0.067 | 17.873 | 0.744 | 20.626 | 1.739 |
| 56 | 0.924 | 0.077 | 20.392 | 0.669 | 22.064 | 2.047 |
| 57 | 0.879 | 0.024 | 20.202 | 0.918 | 22.990 | 1.271 |
| 59 | 0.743 | 0.087 | 20.521 | 1.116 | 27.622 | 3.136 |
| 60 | 0.935 | 0.178 | 18.139 | 0.717 | 19.409 | 3.929 |

| Residue | *R1* | | *R2* | | *R2/R1* | |
|---|---|---|---|---|---|---|
| # | Mean | Error | Mean | Error | Mean | Error |
| 61 | 0.894 | 0.050 | 17.947 | 1.079 | 20.083 | 1.598 |
| 62 | 0.874 | 0.021 | 18.734 | 0.372 | 21.431 | 0.652 |
| 63 | 1.315 | 0.050 | 17.088 | 1.264 | 12.990 | 1.123 |
| 66 | 0.796 | 0.226 | 8.177 | 1.444 | 10.278 | 2.706 |
| 68 | 1.009 | 0.068 | 19.512 | 1.066 | 19.339 | 1.826 |
| 70 | 0.959 | 0.072 | 20.198 | 0.653 | 21.066 | 1.843 |
| 71 | 0.968 | 0.086 | 24.207 | 0.721 | 25.006 | 2.575 |
| 72 | 0.898 | 0.032 | 18.685 | 0.478 | 20.796 | 0.905 |
| 73 | 0.861 | 0.026 | 19.673 | 0.343 | 22.861 | 0.780 |
| 74 | 0.849 | 0.025 | 20.700 | 0.694 | 24.384 | 1.109 |
| 76 | 0.816 | 0.029 | 20.921 | 0.608 | 25.628 | 1.168 |
| 77 | 0.842 | 0.055 | 17.621 | 1.655 | 20.934 | 2.223 |
| 79 | 0.892 | 0.034 | 19.095 | 0.341 | 21.405 | 0.890 |
| 81 | 1.066 | 0.131 | 18.525 | 1.606 | 17.373 | 2.927 |
| 82 | 1.047 | 0.027 | 18.365 | 0.836 | 17.535 | 0.927 |
| 83 | 0.844 | 0.029 | 20.597 | 0.539 | 24.408 | 1.048 |
| 84 | 0.888 | 0.035 | 18.993 | 0.480 | 21.387 | 0.994 |
| 85 | 0.957 | 0.045 | 24.802 | 1.095 | 25.918 | 1.973 |
| 86 | 0.841 | 0.057 | 19.275 | 0.213 | 22.918 | 1.482 |
| 87 | 0.925 | 0.055 | 18.947 | 0.686 | 20.481 | 1.447 |
| 88 | 0.864 | 0.025 | 16.466 | 0.336 | 19.068 | 0.631 |
| 89 | 0.841 | 0.018 | 19.900 | 0.389 | 23.662 | 0.682 |
| 90 | 0.803 | 0.021 | 17.740 | 0.213 | 22.104 | 0.565 |
| 91 | 0.912 | 0.089 | 17.895 | 0.464 | 19.631 | 2.006 |
| 92 | 0.916 | 0.041 | 17.940 | 0.296 | 19.591 | 0.954 |
| 93 | 0.913 | 0.087 | 21.070 | 0.568 | 23.072 | 2.337 |
| 94 | 0.850 | 0.019 | 18.067 | 0.493 | 21.247 | 0.708 |
| 95 | 0.859 | 0.047 | 19.685 | 0.374 | 22.913 | 1.283 |
| 96 | 0.882 | 0.029 | 19.589 | 0.652 | 22.214 | 1.047 |
| 97 | 0.880 | 0.042 | 19.604 | 1.030 | 22.270 | 1.598 |
| 98 | 0.824 | 0.040 | 18.702 | 1.378 | 22.686 | 1.922 |
| 99 | 0.862 | 0.063 | 21.768 | 0.687 | 25.250 | 1.982 |
| 100 | 0.751 | 0.050 | 19.916 | 0.762 | 26.529 | 1.818 |
| 103 | 0.813 | 0.044 | 19.960 | 0.278 | 24.551 | 1.269 |
| 104 | 0.929 | 0.051 | 17.637 | 0.631 | 18.977 | 1.256 |
| 107 | 0.838 | 0.027 | 19.996 | 0.282 | 23.855 | 0.795 |
| 110 | 0.904 | 0.036 | 19.790 | 1.375 | 21.888 | 1.802 |
| 112 | 0.959 | 0.033 | 15.741 | 1.065 | 16.417 | 1.093 |
| 113 | 0.825 | 0.067 | 21.331 | 0.266 | 25.853 | 1.980 |
| 114 | 0.913 | 0.027 | 19.433 | 0.544 | 21.279 | 0.890 |
| 115 | 0.925 | 0.122 | 14.033 | 1.069 | 15.170 | 2.226 |
| 116 | 0.873 | 0.033 | 17.271 | 0.677 | 19.793 | 1.008 |
| 117 | 0.918 | 0.036 | 21.119 | 0.477 | 22.999 | 1.094 |

**Table S3.** $R_1$, $R_2$, and $R_2/R_1$ values for C56S MIF-2

| Residue | *R1* | | *R2* | | *R2/R1* | |
|---|---|---|---|---|---|---|
| # | Mean | Error | Mean | Error | Mean | Error |
| 2 | 1.045 | 0.107 | 17.838 | 1.044 | 17.069 | 2.109 |
| 3 | 0.956 | 0.019 | 18.450 | 0.215 | 19.299 | 0.430 |
| 4 | 0.961 | 0.048 | 17.262 | 0.757 | 17.970 | 1.148 |
| 5 | 0.924 | 0.073 | 17.749 | 0.331 | 19.205 | 1.462 |
| 6 | 0.992 | 0.023 | 18.491 | 0.615 | 18.639 | 0.765 |
| 7 | 0.994 | 0.030 | 17.088 | 0.345 | 17.191 | 0.613 |
| 8 | 0.944 | 0.057 | 19.069 | 0.331 | 20.195 | 1.224 |
| 9 | 0.898 | 0.127 | 16.093 | 1.774 | 17.927 | 2.909 |
| 11 | 0.988 | 0.022 | 17.452 | 0.198 | 17.661 | 0.446 |
| 12 | 1.099 | 0.056 | 17.185 | 2.572 | 15.644 | 2.387 |
| 13 | 0.918 | 0.021 | 17.074 | 0.219 | 18.593 | 0.453 |
| 14 | 0.901 | 0.011 | 15.985 | 0.122 | 17.743 | 0.225 |
| 16 | 0.997 | 0.023 | 16.106 | 0.402 | 16.154 | 0.523 |
| 17 | 0.916 | 0.242 | 8.764 | 2.120 | 9.571 | 2.604 |
| 18 | 1.033 | 0.015 | 18.567 | 0.184 | 17.973 | 0.327 |
| 19 | 1.026 | 0.056 | 16.878 | 0.564 | 16.446 | 1.066 |
| 20 | 0.965 | 0.050 | 17.986 | 0.408 | 18.633 | 1.037 |
| 21 | 0.995 | 0.030 | 17.960 | 0.458 | 18.050 | 0.722 |
| 22 | 0.977 | 0.047 | 18.416 | 0.299 | 18.858 | 0.963 |
| 23 | 0.991 | 0.063 | 17.844 | 0.201 | 18.005 | 1.178 |
| 24 | 0.936 | 0.034 | 17.915 | 0.433 | 19.133 | 0.799 |
| 25 | 1.009 | 0.034 | 17.963 | 0.228 | 17.807 | 0.654 |
| 26 | 0.967 | 0.029 | 17.277 | 0.111 | 17.865 | 0.530 |
| 27 | 0.970 | 0.028 | 17.382 | 0.218 | 17.921 | 0.547 |
| 28 | 0.924 | 0.037 | 15.425 | 0.685 | 16.690 | 0.885 |
| 29 | 0.949 | 0.108 | 17.271 | 0.352 | 18.204 | 2.033 |
| 30 | 0.978 | 0.045 | 16.725 | 0.766 | 17.110 | 1.067 |
| 31 | 0.929 | 0.057 | 18.646 | 0.480 | 20.082 | 1.272 |
| 32 | 0.868 | 0.039 | 18.225 | 0.088 | 20.995 | 0.834 |
| 34 | 0.969 | 0.019 | 21.925 | 0.702 | 22.627 | 0.974 |
| 35 | 0.891 | 0.046 | 24.201 | 0.978 | 27.154 | 1.928 |
| 36 | 0.933 | 0.106 | 17.940 | 0.550 | 19.232 | 2.159 |
| 37 | 0.981 | 0.027 | 16.581 | 0.451 | 16.896 | 0.631 |
| 38 | 0.572 | 0.173 | 26.261 | 4.269 | 45.877 | 13.459 |
| 39 | 0.991 | 0.052 | 17.232 | 0.442 | 17.388 | 1.020 |
| 40 | 0.976 | 0.062 | 16.938 | 0.557 | 17.361 | 1.225 |
| 41 | 0.927 | 0.020 | 18.012 | 0.331 | 19.434 | 0.527 |
| 42 | 0.958 | 0.059 | 18.106 | 0.856 | 18.903 | 1.444 |
| 44 | 0.847 | 0.049 | 17.841 | 0.735 | 21.053 | 1.353 |
| 45 | 0.864 | 0.042 | 16.631 | 0.548 | 19.258 | 1.001 |
| 46 | 0.975 | 0.028 | 18.047 | 0.280 | 18.517 | 0.605 |
| 47 | 0.943 | 0.037 | 16.537 | 0.675 | 17.546 | 0.935 |
| 49 | 0.983 | 0.025 | 17.117 | 0.325 | 17.408 | 0.548 |
| 50 | 0.980 | 0.043 | 18.235 | 0.579 | 18.600 | 1.000 |
| 51 | 1.044 | 0.036 | 16.260 | 0.381 | 15.576 | 0.659 |
| 52 | 1.069 | 0.015 | 15.684 | 0.196 | 14.675 | 0.280 |
| 54 | 1.058 | 0.010 | 18.064 | 0.044 | 17.074 | 0.172 |
| 57 | 0.989 | 0.107 | 23.392 | 7.277 | 23.649 | 9.811 |
| 59 | 1.013 | 0.084 | 18.825 | 0.581 | 18.577 | 1.689 |
| 60 | 1.017 | 0.051 | 17.581 | 0.275 | 17.284 | 0.938 |
| 61 | 0.983 | 0.026 | 18.577 | 0.745 | 18.893 | 0.925 |
| 62 | 0.997 | 0.028 | 17.957 | 0.435 | 18.010 | 0.671 |
| 63 | 0.951 | 0.075 | 17.337 | 0.956 | 18.221 | 1.682 |
| 66 | 1.026 | 0.144 | 14.671 | 0.682 | 14.305 | 2.166 |
| 69 | 1.070 | 0.133 | 27.278 | 1.295 | 25.494 | 3.896 |
| 70 | 1.073 | 0.092 | 15.662 | 0.687 | 14.603 | 1.472 |
| 71 | 1.058 | 0.045 | 19.091 | 0.744 | 18.049 | 1.108 |
| 72 | 1.033 | 0.037 | 17.643 | 0.102 | 17.073 | 0.658 |
| 73 | 1.058 | 0.018 | 18.106 | 0.341 | 17.118 | 0.449 |
| 74 | 1.025 | 0.049 | 21.829 | 1.868 | 21.297 | 2.431 |
| 76 | 1.034 | 0.020 | 19.331 | 0.523 | 18.693 | 0.659 |
| 77 | 0.952 | 0.081 | 18.103 | 1.134 | 19.008 | 1.960 |
| 79 | 0.978 | 0.025 | 17.212 | 0.169 | 17.590 | 0.473 |
| 80 | 0.954 | 0.082 | 18.804 | 1.022 | 19.707 | 1.979 |
| 81 | 0.993 | 0.056 | 19.257 | 0.400 | 19.391 | 1.187 |
| 82 | 0.971 | 0.034 | 14.476 | 0.268 | 14.910 | 0.563 |
| 83 | 0.933 | 0.041 | 18.386 | 0.385 | 19.710 | 0.927 |
| 84 | 1.023 | 0.029 | 18.165 | 0.227 | 17.751 | 0.576 |
| 85 | 0.994 | 0.040 | 18.580 | 1.056 | 18.692 | 1.322 |
| 86 | 0.903 | 0.031 | 17.062 | 0.241 | 18.888 | 0.639 |
| 87 | 0.911 | 0.052 | 16.941 | 0.275 | 18.601 | 1.026 |
| 88 | 0.927 | 0.037 | 16.589 | 0.240 | 17.900 | 0.722 |
| 89 | 0.954 | 0.011 | 18.146 | 0.286 | 19.017 | 0.365 |
| 90 | 0.838 | 0.031 | 16.072 | 0.731 | 19.174 | 0.983 |
| 91 | 0.903 | 0.041 | 16.722 | 0.224 | 18.512 | 0.811 |
| 92 | 1.032 | 0.049 | 17.746 | 0.948 | 17.191 | 1.243 |
| 93 | 1.044 | 0.024 | 20.329 | 0.517 | 19.463 | 0.731 |
| 94 | 1.027 | 0.009 | 17.259 | 0.192 | 16.812 | 0.240 |
| 95 | 0.994 | 0.023 | 17.286 | 0.149 | 17.390 | 0.430 |
| 96 | 0.973 | 0.016 | 17.581 | 0.464 | 18.073 | 0.542 |
| 97 | 0.980 | 0.039 | 21.263 | 1.008 | 21.688 | 1.477 |
| 98 | 1.007 | 0.025 | 16.554 | 0.444 | 16.441 | 0.582 |
| 99 | 1.057 | 0.069 | 18.100 | 0.472 | 17.128 | 1.276 |
| 100 | 0.969 | 0.043 | 17.892 | 0.375 | 18.465 | 0.895 |
| 103 | 0.859 | 0.108 | 20.475 | 0.935 | 23.833 | 2.904 |
| 104 | 1.087 | 0.038 | 15.944 | 0.353 | 14.667 | 0.637 |
| 106 | 0.992 | 0.095 | 18.911 | 1.484 | 19.062 | 2.408 |
| 107 | 1.008 | 0.012 | 19.011 | 0.086 | 18.857 | 0.241 |
| 110 | 0.938 | 0.046 | 20.044 | 0.924 | 21.367 | 1.482 |
| 113 | 1.078 | 0.060 | 19.736 | 0.335 | 18.311 | 1.175 |
| 114 | 1.022 | 0.014 | 17.289 | 0.132 | 16.917 | 0.275 |
| 116 | 0.963 | 0.073 | 21.786 | 1.301 | 22.614 | 2.331 |
| 117 | 1.074 | 0.027 | 18.965 | 0.175 | 17.652 | 0.514 |

**Table S4.** $R_1$, $R_2$, and $R_2/R_1$ values for C56S MIF-2 bound to Ebselen

| Residue | *R1* | | *R2* | | *R2/R1* | | Residue | *R1* | | *R2* | | *R2/R1* | |
|---|---|---|---|---|---|---|---|---|---|---|---|---|---|
| # | Mean | Error | Mean | Error | Mean | Error | # | Mean | Error | Mean | Error | Mean | Error |
| 2 | 0.911 | 0.003 | 20.133 | 0.196 | 22.106 | 0.242 | 62 | 0.949 | 0.040 | 19.048 | 0.768 | 20.076 | 1.188 |
| 3 | 0.883 | 0.048 | 16.507 | 0.346 | 18.703 | 1.017 | 63 | 0.926 | 0.011 | 15.916 | 0.537 | 17.189 | 0.533 |
| 4 | 0.842 | 0.015 | 16.941 | 0.077 | 20.125 | 0.323 | 66 | 1.754 | 0.098 | 13.441 | 1.194 | 7.663 | 0.939 |
| 5 | 0.881 | 0.027 | 20.239 | 0.496 | 22.971 | 0.896 | 70 | 0.962 | 0.055 | 20.551 | 0.386 | 21.373 | 1.341 |
| 6 | 0.853 | 0.013 | 17.986 | 0.329 | 21.079 | 0.472 | 71 | 0.974 | 0.048 | 20.725 | 0.143 | 21.285 | 1.099 |
| 7 | 0.867 | 0.021 | 18.188 | 0.104 | 20.971 | 0.478 | 72 | 0.907 | 0.013 | 17.979 | 0.248 | 19.831 | 0.385 |
| 8 | 0.911 | 0.031 | 19.260 | 0.259 | 21.148 | 0.770 | 73 | 0.913 | 0.039 | 19.369 | 0.312 | 21.209 | 0.957 |
| 9 | 0.909 | 0.032 | 17.809 | 0.251 | 19.590 | 0.711 | 74 | 0.918 | 0.035 | 17.999 | 1.619 | 19.600 | 1.849 |
| 11 | 0.851 | 0.044 | 18.723 | 1.069 | 22.000 | 1.633 | 76 | 0.932 | 0.016 | 21.492 | 1.735 | 23.060 | 2.177 |
| 12 | 0.880 | 0.022 | 15.886 | 0.725 | 18.046 | 0.819 | 77 | 0.867 | 0.040 | 16.442 | 0.229 | 18.974 | 0.840 |
| 13 | 0.866 | 0.033 | 18.488 | 0.099 | 21.353 | 0.765 | 79 | 0.920 | 0.012 | 18.646 | 0.450 | 20.269 | 0.551 |
| 14 | 0.911 | 0.021 | 19.008 | 0.120 | 20.871 | 0.478 | 80 | 0.959 | 0.086 | 19.470 | 0.681 | 20.308 | 2.008 |
| 16 | 0.875 | 0.045 | 16.033 | 0.113 | 18.326 | 0.885 | 81 | 0.906 | 0.035 | 19.212 | 0.364 | 21.210 | 0.896 |
| 18 | 0.978 | 0.052 | 20.141 | 0.547 | 20.604 | 1.288 | 82 | 0.935 | 0.021 | 19.186 | 0.169 | 20.530 | 0.493 |
| 19 | 0.871 | 0.028 | 19.470 | 0.192 | 22.352 | 0.697 | 83 | 0.867 | 0.031 | 19.011 | 0.222 | 21.939 | 0.783 |
| 20 | 1.168 | 0.027 | 11.663 | 14.300 | 9.984 | 7.649 | 84 | 0.946 | 0.023 | 18.657 | 0.136 | 19.720 | 0.502 |
| 21 | 0.931 | 0.025 | 20.080 | 0.373 | 21.566 | 0.717 | 85 | 0.948 | 0.025 | 24.716 | 0.412 | 26.075 | 0.903 |
| 22 | 0.905 | 0.017 | 20.496 | 1.138 | 22.648 | 1.442 | 86 | 0.866 | 0.021 | 18.896 | 0.092 | 21.825 | 0.495 |
| 23 | 0.814 | 0.083 | 15.328 | 1.727 | 18.838 | 2.411 | 87 | 0.921 | 0.031 | 18.103 | 1.127 | 19.660 | 1.350 |
| 25 | 1.052 | 0.026 | 17.992 | 0.090 | 17.100 | 0.481 | 88 | 0.854 | 0.034 | 16.923 | 0.897 | 19.817 | 1.197 |
| 26 | 0.893 | 0.025 | 19.387 | 0.470 | 21.714 | 0.796 | 89 | 0.887 | 0.034 | 19.704 | 0.260 | 22.207 | 0.870 |
| 27 | 0.903 | 0.072 | 19.157 | 8.062 | 21.207 | 9.299 | 90 | 0.855 | 0.017 | 16.565 | 0.262 | 19.364 | 0.445 |
| 28 | 0.850 | 0.021 | 17.334 | 0.150 | 20.402 | 0.494 | 91 | 0.874 | 0.020 | 18.481 | 0.321 | 21.142 | 0.586 |
| 29 | 0.971 | 0.046 | 20.454 | 0.354 | 21.068 | 1.101 | 92 | 0.917 | 0.030 | 19.608 | 0.175 | 21.392 | 0.722 |
| 30 | 0.909 | 0.032 | 17.065 | 0.197 | 18.771 | 0.666 | 93 | 0.923 | 0.063 | 20.004 | 0.178 | 21.684 | 1.467 |
| 31 | 0.829 | 0.037 | 19.600 | 0.173 | 23.657 | 0.953 | 94 | 0.907 | 0.035 | 19.635 | 0.117 | 21.638 | 0.810 |
| 32 | 1.269 | 0.030 | 22.031 | 0.367 | 17.365 | 0.646 | 95 | 0.890 | 0.023 | 18.413 | 0.106 | 20.678 | 0.522 |
| 34 | 0.902 | 0.030 | 25.648 | 1.589 | 28.443 | 2.581 | 96 | 0.708 | 0.032 | 8.104 | 0.433 | 11.451 | 0.477 |
| 35 | 0.870 | 0.010 | 21.053 | 0.175 | 24.211 | 0.346 | 97 | 0.896 | 0.034 | 21.053 | 0.447 | 23.495 | 1.029 |
| 36 | 0.928 | 0.043 | 10.516 | 2.718 | 11.337 | 1.730 | 98 | 0.858 | 0.025 | 17.787 | 0.280 | 20.722 | 0.630 |
| 38 | 0.818 | 0.033 | 16.226 | 0.393 | 19.844 | 0.806 | 99 | 0.846 | 0.041 | 16.966 | 0.251 | 20.054 | 0.929 |
| 39 | 0.921 | 0.050 | 19.357 | 0.171 | 21.022 | 1.142 | 100 | 0.844 | 0.030 | 18.688 | 0.202 | 22.145 | 0.739 |
| 40 | 0.933 | 0.017 | 17.816 | 0.079 | 19.099 | 0.359 | 103 | 0.929 | 0.046 | 19.283 | 0.248 | 20.748 | 1.049 |
| 41 | 0.865 | 0.028 | 17.995 | 0.270 | 20.803 | 0.692 | 104 | 0.898 | 0.023 | 17.209 | 0.315 | 19.153 | 0.577 |
| 42 | 0.987 | 0.014 | 18.262 | 0.166 | 18.499 | 0.317 | 106 | 1.058 | 0.034 | 20.812 | 1.024 | 19.665 | 1.290 |
| 44 | 0.869 | 0.029 | 17.771 | 0.382 | 20.455 | 0.756 | 107 | 0.903 | 0.029 | 19.073 | 0.238 | 21.114 | 0.708 |
| 46 | 0.870 | 0.046 | 20.799 | 1.490 | 23.898 | 2.239 | 110 | 0.889 | 0.090 | 19.088 | 0.912 | 21.474 | 2.313 |
| 47 | 0.849 | 0.031 | 17.091 | 0.128 | 20.133 | 0.683 | 111 | 1.020 | 0.047 | 32.010 | 1.155 | 31.389 | 2.510 |
| 49 | 0.952 | 0.060 | 17.630 | 0.301 | 18.512 | 1.222 | 112 | 0.942 | 0.045 | 18.440 | 0.869 | 19.583 | 1.307 |
| 51 | 1.022 | 0.027 | 17.637 | 0.286 | 17.263 | 0.570 | 113 | 0.930 | 0.022 | 20.350 | 0.826 | 21.876 | 1.099 |
| 54 | 2.034 | 0.035 | 2.076 | 0.170 | 1.021 | 0.040 | 114 | 0.931 | 0.021 | 18.546 | 0.090 | 19.918 | 0.464 |
| 57 | 0.842 | 0.030 | 26.539 | 0.378 | 31.529 | 1.204 | 115 | 1.199 | 0.073 | 24.492 | 1.809 | 20.424 | 2.538 |
| 59 | 0.888 | 0.032 | 19.084 | 0.647 | 21.489 | 1.052 | 116 | 0.936 | 0.049 | 23.458 | 1.597 | 25.053 | 2.518 |
| 60 | 0.968 | 0.050 | 18.212 | 0.250 | 18.813 | 1.027 | 117 | 0.966 | 0.032 | 20.492 | 0.183 | 21.209 | 0.757 |
| 61 | 0.835 | 0.023 | 18.162 | 0.238 | 21.740 | 0.597 | | | | | | | |
